# Regulation strategies for two-output biomolecular networks

**DOI:** 10.1101/2022.02.28.482258

**Authors:** Emmanouil Alexis, Carolin CM Schulte, Luca Cardelli, Antonis Papachristodoulou

## Abstract

Feedback control theory facilitates the development of self-regulating systems with desired performance which are predictable and insensitive to disturbances. Feedback regulatory topologies are found in many natural systems and have been of key importance in the design of reliable synthetic bio-devices operating in complex biological environments. Here, we study control schemes for biomolecular processes with two outputs of interest, expanding previous traditional concepts describing one-output systems. Regulation of such processes may unlock new design possibilities but can be challenging due to coupling interactions; also potential disturbances applied on one of the outputs may affect both. We therefore propose architectures for robustly manipulating the ratio and linear combinations of the outputs as well as each of the outputs independently. To demonstrate their characteristics, we apply these architectures to a simple process of two mutually activated biomolecular species. We also highlight the potential for experimental implementation by exploring synthetic realizations both *in vivo* and *in vitro*. This work presents an important step forward in building bio-devices capable of sophisticated functions.

## 1 Introduction

For more than two decades we have witnessed significant advances in the highly interdisciplinary field of synthetic biology whose goal it is to harness engineering approaches in order to realize genetic networks that produce user-defined cellular outcomes. These advances have the potential to transform several aspects of our life by providing efficient solutions to a long list of critical global issues related to food security, healthcare, energy and the environment [1–6]. A fundamental characteristic of living systems is the presence of multi-scale feedback mechanisms facilitating their functioning and survival [7, 8]. Feedback control enables a self-regulating system to adjust its current and future actions by sensing the state of its outputs. This seems to be the answer to a number of major challenges that prevent successful implementation of synthetic genetic circuits and keep innovative endeavours in the field trapped at a laboratory stage. Control theory offers a rich toolkit of powerful techniques to design and manipulate biological systems and enable the reliable function of next-generation synthetic biology applications [9–13].

Engineering life aims at constructing modular biomolecular devices which are able to operate in a controllable and predictable way in constantly changing environments with a high level of burden and cross-talk. It is therefore a requirement for them to be resilient to context-dependent effects and adapt to external environmental perturbations. Several control approaches inspired by both natural and technological systems have recently been proposed allowing for effective and robust regulation of biological networks *in vivo* and/or *in vitro* [14–20]. Despite some conceptual differences, all of these studies focus on biomolecular systems with one output of interest, such as the expression of a single protein.

Building advanced bio-devices capable of performing more sophisticated computations and tasks requires the design of genetic circuits where multiple inputs are applied and multiple outputs are measured. In control engineering these types of systems are also known as multi-input multi-output or MIMO systems [21]. This may be the key for achieving control of the whole cell, which can be regarded as a very complex MIMO bio-device itself. Regulation of processes comprising multiple interacting variables of interest can be challenging since there may be interactions between inputs and outputs. Thus, a change in any input may affect all outputs. At the same time, attempts to apply feedback control by “closing the loop” could be impaired by input - output pairing. Addressing such problems therefore requires alternative, suitably adjusted regulation schemes which take into account the presence of mutual internal interactions in the network to be controlled (open-loop system). The research area of MIMO control bio-systems has up until now remained relatively unexplored. There have been only a few studies towards this direction coming mainly from the field of cybergenetics where a computer is a necessary part of the control feedback loop [22, 23]. In contrast, substantial progress has been made in a closely related area, namely MIMO logic bio-circuits which are able to realize Boolean functions [24, 25] while “multi-layer” control concepts for one-output processes [26, 27] and resource allocation in gene expression [28] have also been proposed.

In this paper, we investigate regulation strategies for biomolecular networks with two outputs of interest which can correspond, for example, to the concentration of two different proteins inside the cell, assuming the presence of mutual interactions. Both the open-loop and the closed-loop system (open-loop system within a feedback control configuration) are represented by chemical reaction networks (CRNs) obeying the law of mass action [8]. Consequently, the entire regulation process takes place in the biological context of interest without the use of computer-aided methods. We exploit “multi-loop” concepts based on two independent feedback loops as well as concepts where the control action is carried out jointly considering both outputs simultaneously. Moreover, our designs take advantage of the adaptation benefits stemming from integral feedback action realized through molecular sequestration [29].

Specifically, we present regulating architectures, which we refer to as regulators, capable of achieving one of the following control objectives: robustly driving a) the ratio of the outputs; b) a linear combination of the outputs; and c) each of the outputs to a desired value (set point). At steady state, the architectures of a) and b) result in two coupled outputs which can still affect each other, albeit in a specific way dictated by the respective control approach. On the other hand, the architectures for c) achieve steady state decoupling, thus making the two outputs independent of each other. Our control schemes can be used for regulation of any arbitrary open-loop process provided that the resulting closed-loop system has a finite, positive steady state and the closed-loop system converges to that steady state as time goes to infinity (closed-loop (asymptotic) stability). Thus, the present analysis focuses exclusively on such scenarios. Furthermore, we mathematically and computationally demonstrate their special characteristics by applying these schemes to a simple biological process of two mutually activating species. Finally, to highlight their biological relevance and motivate further experimental investigation, we explore potential implementations of our designs.

## Results

### 2 Control schemes with steady state coupling

In Figure 1A we show a general biomolecular process with two outputs of interest for which we first present two bio-controllers aiming to regulate the ratio and an arbitrary linear combination of the outputs, respectively. The different types of biomolecular reactions as well as their graphical representations used in this work are presented in Figure 1B.

**Figure 1:**
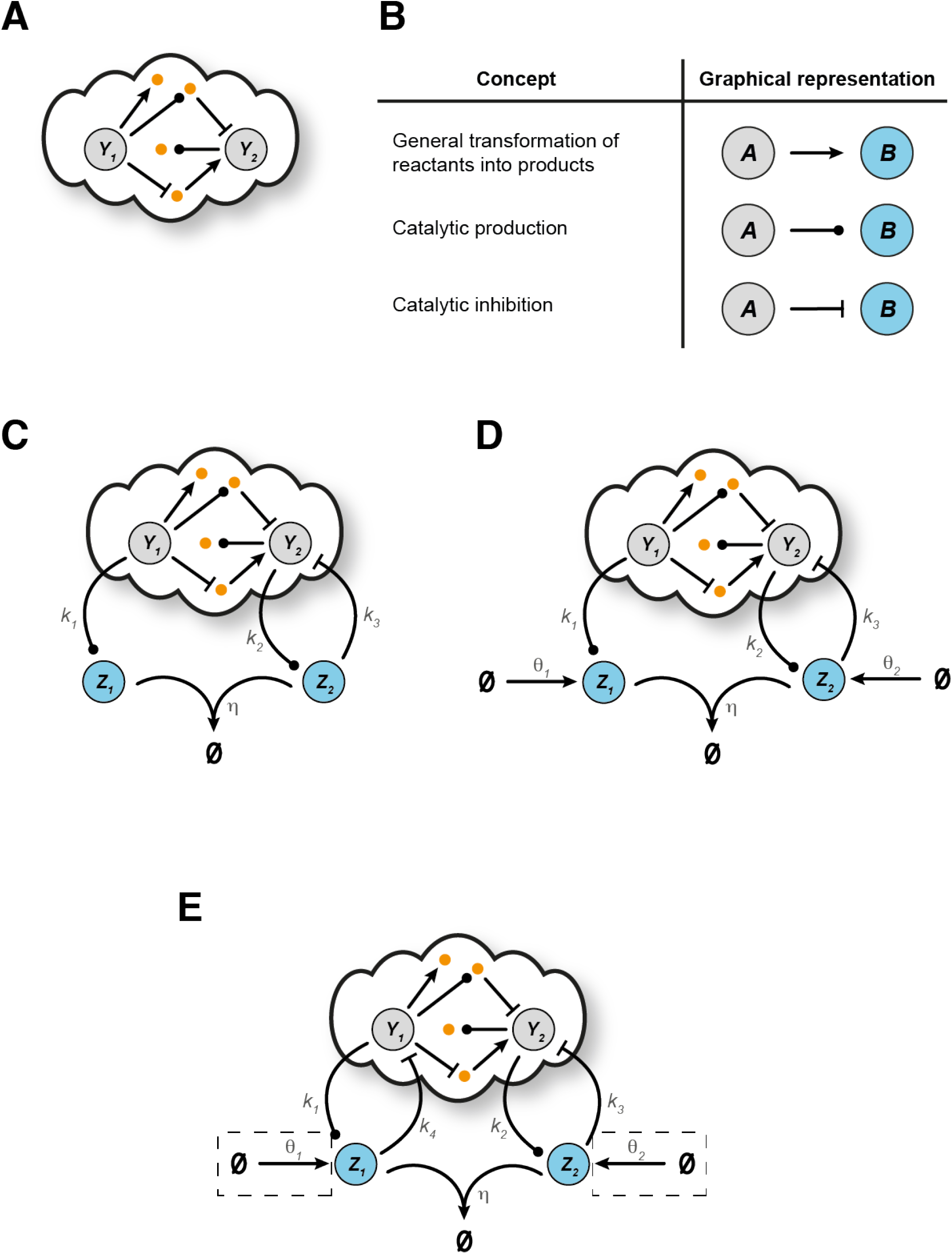
Open-loop biomolecular network and control architectures with steady state coupling. **A** Schematic representation of a general biomolecular network with two output species of interest, *Y*_1_, *Y*_2_, and an arbitrary number of other species and/or biomolecular interactions. **B** Graphical representation of the different types of biochemical reactions adopted from our previous work [37]. Schematic representation of a general closed-loop architecture using **C** R-Regulator (CRN (1)), **D** LC-Regulator (CRN (5)), **E** R- and LC-Regulator with an additional inhibitory reaction (the biological parts enclosed in dashed boxes are only required for LC-Regulator).

#### 2.1 Regulating the ratio of outputs

Figure 1C illustrates a motif which we call R-Regulator and consists of the following reactions:

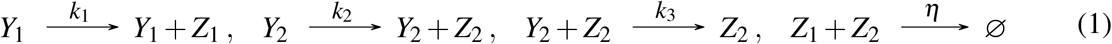

This controller consists of two species, *Z*_1_ and *Z*_2_, which annihilate each other. The production of *Z*_1_,*Z*_2_ is catalyzed by the target species *Y*_1_, *Y*_2_, respectively while *Y*_2_ is also inhibited by *Z*_2_.

The dynamics of the R-Regulator are described by the following system of Ordinary Differential Equations (ODEs):

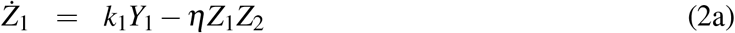

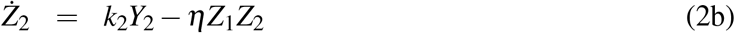

Equations (2a)-(2b) give rise to a non physical memory variable which enables integration, i.e.:

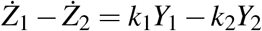

or

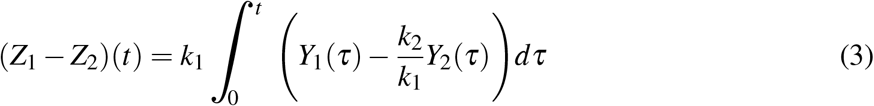

As a result, assuming closed-loop stability (*Ż*_1_, *Ż*_2_ → 0 as t → ∞), we get:

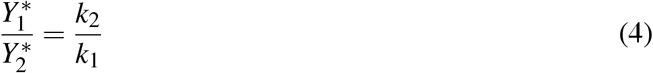

where the ^*∗*^ notation indicates the steady state concentration of a species. As can be seen, the integrand in Equation (3) corresponds to an error quantity which converges to zero over time, thus guaranteeing that the output ratio 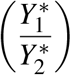 will converge to the set point 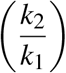. It is important to note that the aforementioned stability depends on the structure of the open-loop process, which is unknown here, as well as the set of the reaction rates/parameter values we select for the closed-loop system.

#### 2.2 Regulating a linear combination of the outputs

In Figure 1D a second motif, which we call LC-Regulator, is depicted. The only difference to the R-Regulator is that species *Z*_1_, *Z*_2_ are also produced through two independent processes with constant rates *θ*_1_, *θ*_2_, respectively. More analytically, the corresponding reaction network is:

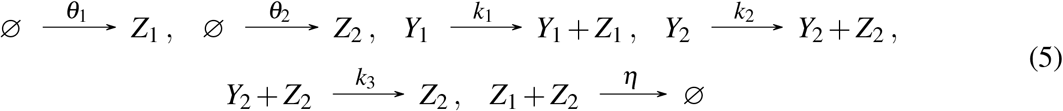

The dynamics of LC-Regulator is given by the set of ODEs:

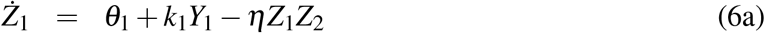

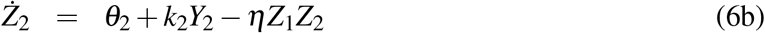

Similar to before, in order to see the memory function involved, we subtract Equations (6a) - (6b) and integrate to get:

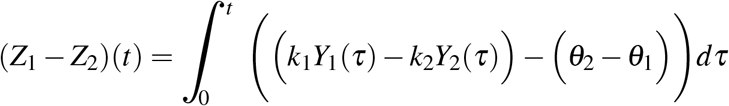

Under the assumption of closed-loop stability (*Ż*_1_, *Ż*_2_ → 0 as t → ∞), we have at steady state:

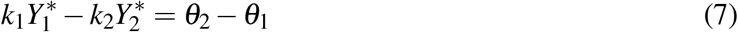

An interesting feature of LC-Regulator is that Equation (7) can be adjusted as desired by modifying the production reactions regarding *Z*_1_, *Z*_2_. Imagine, for example, replacing 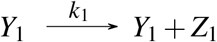 with 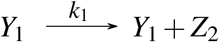 in CRN (5). Then the term *k*_1_*Y*_1_ would appear in Equation (6b) instead of (6a) resulting in 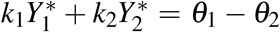 at steady state. Note that in this work we focus on CRN (5) presented initially as the representative of LC-Regulator.

Finally, in Figure 1E we show an alternative version of the controllers presented above. Specifically, the inhibitory reaction 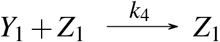 has been added to the R- or LC-Regulator in this scenario. Note that this additional reaction does not change the dynamics of the controllers - Equations (2a)- (2b) and (6a) - (6b) still hold for R-Regulator and LC-Regulator, respectively. Despite the increase in complexity, the additional reaction strengthens the regulatory ability of the controllers in the sense that control action is now applied on both target species. This could, for example, be useful to make closed-loop stability more robust. These slightly modified motifs are further discussed in Section **Closed-loop stability**.

### 3 Control schemes with steady state decoupling

We now present three alternative bio-controllers, which we call D-Regulator I, II and III, capable of achieving independent control of each output in the arbitrary biomolecular process (Figure 1A). In particular, D-Regulators are able to drive each output species to a desired steady state concentration unaffected by the behaviour of the other species.

#### 3.1 D-Regulator I

The set of reactions describing D-Regulator I (Figure 2A) is:

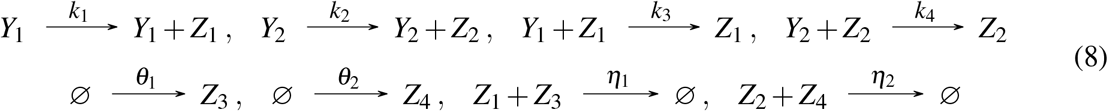

**Figure 2:**
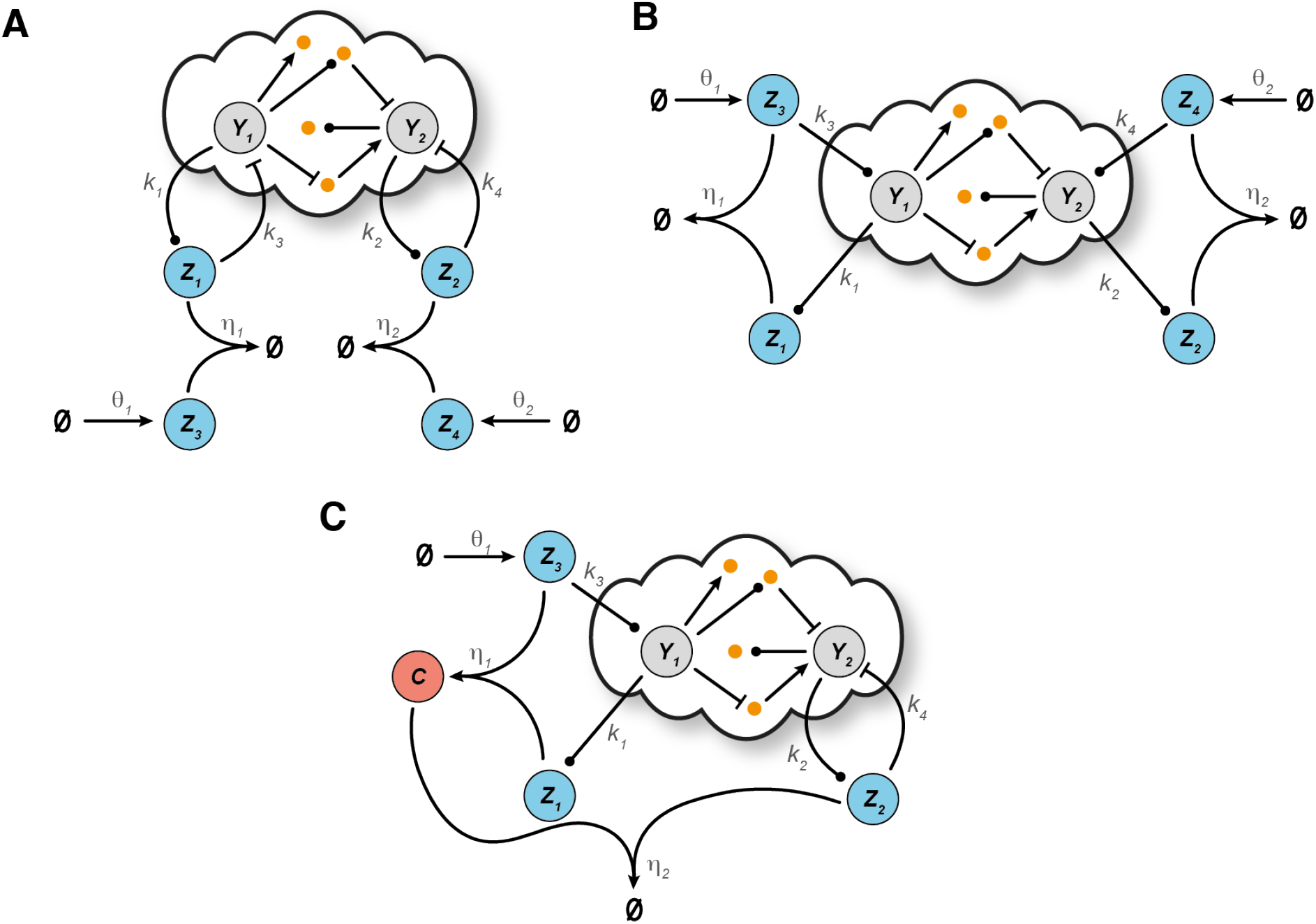
Control architectures with steady state decoupling. Schematic representation of a general closed-loop architecture using **A** D-Regulator I (CRN (8)), **B** D-Regulator II (CRN (12)) and **C** D-Regulator III (CRN (13)).

Here there are four controller species. The target species *Y*_1_, *Y*_2_ catalyze the formation of two of them, *Z*_1_, *Z*_2_, which, in turn, inhibit the former. In addition, *Z*_3_, *Z*_4_, which are produced independently at a constant rate, participate in annihilation reactions with *Z*_1_ and *Z*_2_, respectively.

The dynamics of D-Regulator I can be modelled using the following set of ODEs:

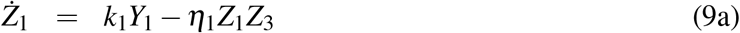

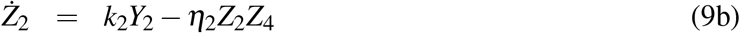

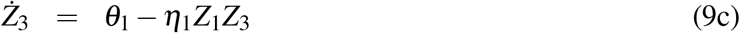

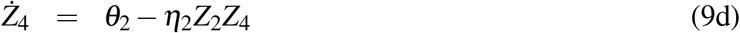

In contrast to the regulation strategies presented in the preceding section, D-Regulator I includes two memory variables which carry out integral action independently. Indeed, combining Equations (9a), (9c) results in:

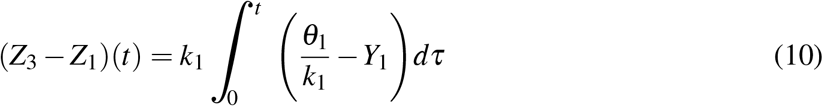

while combining Equations (9b), (9d) gives:

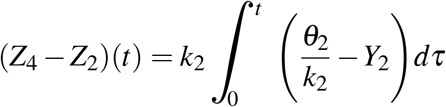

Consequently, the steady state output concentrations under the assumption of closed-loop stability (*Ż*_1_, *Ż*_2_ → 0 as t → ∞) are:

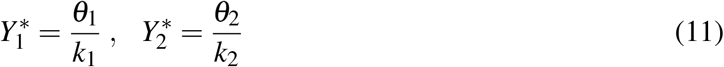

#### 3.2 D-Regulator II

By using four controller species as before and exploiting the control concept introduced in [29], we construct D-Regulator II (Figure 2B) consisting of the following reactions:

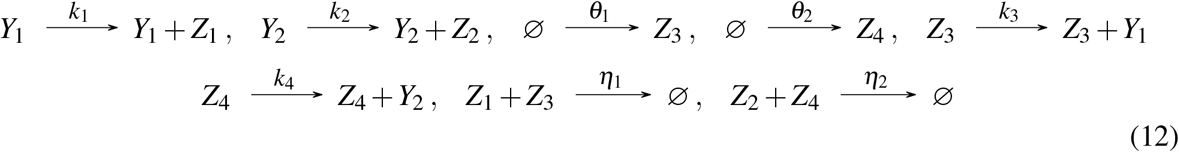

In this case, species *Z*_3_, *Z*_4_ catalyze the formation of the target species *Y*_1_, *Y*_2_, respectively, and *Z*_3_, *Z*_4_ are produced at a constant rate. Furthermore, species *Z*_1_, *Z*_2_ are catalytically produced by *Y*_1_, *Y*_2_, respectively, while the pairs *Z*_1_-*Z*_3_ and *Z*_2_-*Z*_4_ participate in an annihilation reaction.

Note that the species of D-Regulator II are described by the same ODE model as D-Regulator I (Equations (9a)-(9d)). Thus, the memory variables involved as well as the steady state output behaviour (Equation (18)) are identical in these two motifs (provided that close-loop stability is guaranteed). Nonetheless, in general, regulating the same open-loop process via the aforementioned controllers results in different output behaviour until an equilibrium is reached or, in other words, we have different transient responses. This is because of the different topological characteristics of the two motifs which cannot be captured by focusing only on the controller dynamics: considering closed-loop dynamics is required, which is addressed in a later section.

#### 3.3 D-Regulator III

The last bio-controller presented in this study is D-Regulator III (Figure 2C) whose structure is composed of the following reactions:

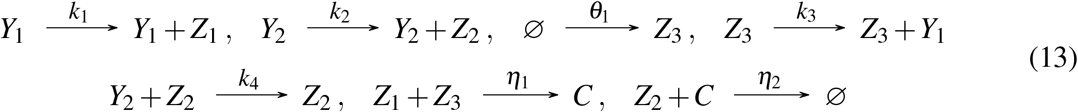

Here there are three controller species. *Z*_1_, *Z*_3_ interact with the target species *Y*_1_ as well as with each other in the same way as in D-Regulator II. The complex *C*, which is formed by the binding of *Z*_1_, *Z*_3_, and the third controller species, *Z*_2_, can annihilate each other. Finally, the target species *Y*_2_ catalyzes the production of *Z*_2_ which, in turn, inhibits *Y*_2_ analogous to D-Regulator I.

The dynamics of D-Regulator III can be described by the following set of ODEs:

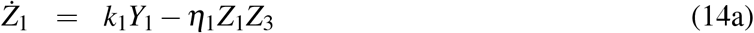

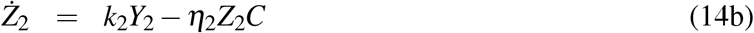

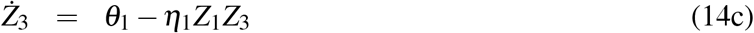

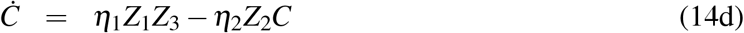

Similar to the other D-Regulators, the memory function responsible for the regulation of the output *Y*_1_ is carried out by the (non-physical) quantity *Z*_3_ −*Z*_1_ (Equation (10)). However, the memory variable related to the output *Y*_2_ is realized in a different way than before. More specifically, combining Equations (14b)-(14d) yields:

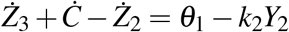

or

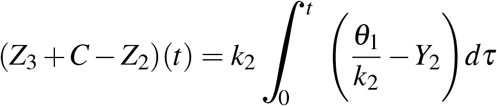

Therefore, assuming closed-loop stability, i.e. *Ż*_1_, *Ż*_2_ → 0 as t → ∞, the steady state output behaviour is:

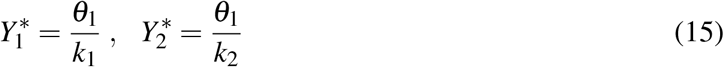

### 4 Specifying the biological network to be controlled

We now turn our focus to a specific two-output open-loop network which will henceforward take the place of the abstract “cloud” process in the preceding sections. This will allow us to implement *in silico* the proposed control motifs and demonstrate the properties discussed above (see **Implementing the proposed regulation strategies**). In addition, we will explore potential experimental realizations of the resulting closed-loop networks (see **Experimental realization**).

Figure 3A illustrates a simple biological network comprised of two general birth-death processes involving two target species, *Y*_1_, *Y*_2_. These species are coupled in the sense that each of them is able to catalyze the formation of the other. Such motifs of positive feedback action are ubiquitous in biological systems [30–32]. In particular, we have the reactions:

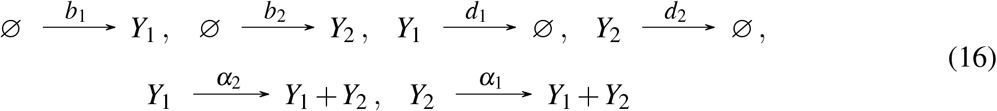

**Figure 3:**
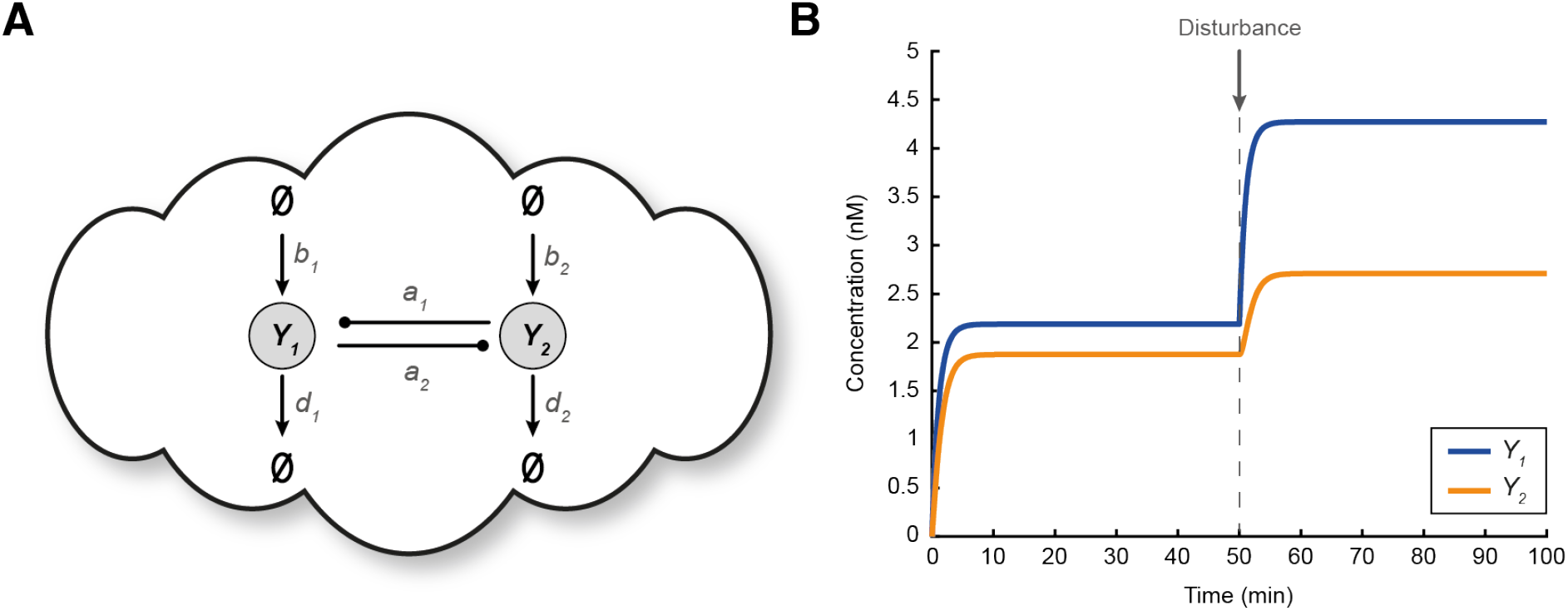
Specifying the open-loop biomolecular network. **A** A simple biological process with two mutually activating output species *Y*_1_, *Y*_2_, described by CRN (16). **B** Simulated response of the topology in **A** using the ODE model (17) with the following parameters: *b*_1_ = 2 nM min^−1^, *b*_2_ = 1 nM min^−1^, *d*_1_ = *d*_2_ = 1 min^−1^, *α*_1_ = 0.1 min^−1^, *α*_2_ = 0.4 min^−1^. At time *t* = 50 min, a disturbance on *Y*_1_ is introduced which affects both output species. More specifically, the value of parameter *b*_1_ changes from 2 to 4.

which can be modelled as:

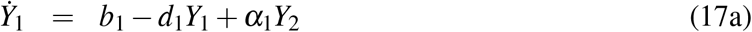

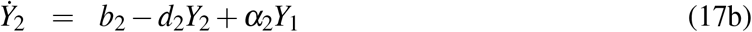

For any *d*_1_*d*_2_ > *α*_1_*α*_2_, ODE system (17a)-(17b) has the following unique positive steady state:

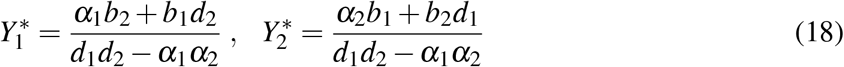

which is (globally) exponentially stable (see Section S2 of the supplementary material).

Note that for this system, a change in any of the reaction rates of network (16) due to, for instance, undesired disturbances, will affect the behaviour of both species *Y*_1_ and *Y*_2_ (Figure 3B).

### 5 Implementing the proposed regulation strategies

We now demonstrate the efficiency of the bio-controllers introduced in **Control schemes with steady state coupling** and **Control schemes with steady state decoupling** by regulating the open-loop network (16) presented in **Specifying the biological network to be controlled**. A detailed analysis of the steady state behaviour of the resulting closed-loop processes can be found in Section S3 of the supplementary material.

We show in Figure 4 that R-Regulator and LC-Regulator are capable of driving the ratio and a desired linear combination of the output species to the set point of our choice in the presence of constant disturbances, respectively. Similarly, we illustrate in Figure 5 the ability of D-Regulators to robustly steer each of the output species towards a desired value independently, thus cancelling the steady state coupling.

**Figure 4:**
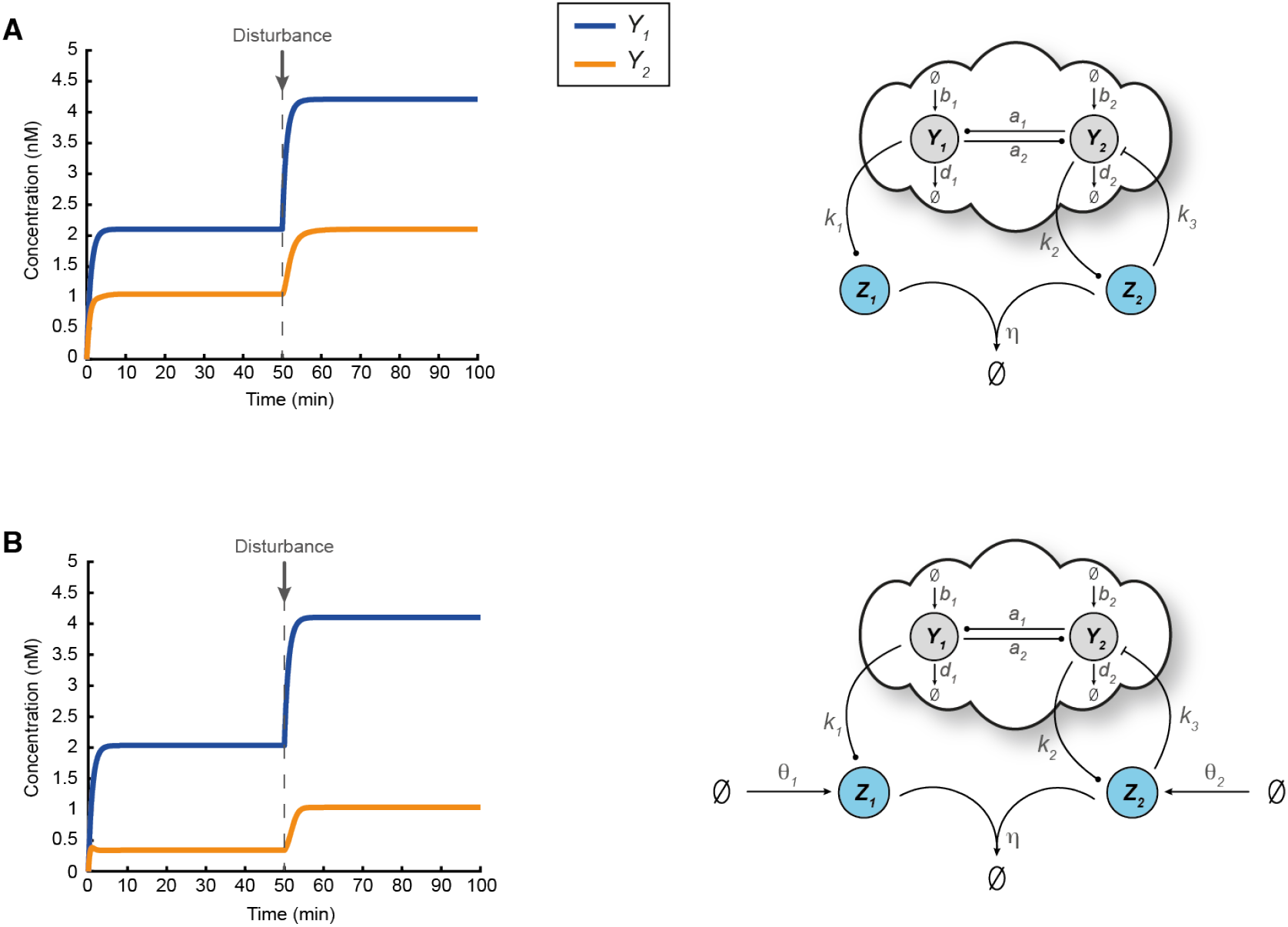
Regulating the ratio and an arbitrary linear combination of the outputs. **A** A closed-loop architecture based on the open-loop network shown in Figure 3A and R-Regulator. For the simulated response presented here the following parameters are used: *k*_1_ = 0.5 min^−1^, *k*_2_ = 1 min^−1^, *k*_3_ = 2 nM^−1^ min^−1^, *η* = 10 nM^−1^ min^−1^ while the rest of the parameters (associated with the open-loop network) are the same as the ones used in Figure 3B. At time *t* = 50 min, a disturbance is applied (same as in Figure 3B) which alters the output steady states. Nevertheless, 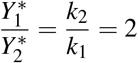 always holds (Equation (4)). **B** A closed-loop architecture based on the open-loop network shown in Figure 3A and LC-Regulator. For the simulated response presented here the following parameters are used: *k*_1_ = 1 min^−1^, *k*_2_ = 3 min^−1^, *k*_3_ = 2 nM^−1^ min^−1^, *η* = 10 nM^−1^ min^−1^, *θ*_1_ = 4 nM min^−1^, *θ*_2_ = 5 nM min^−1^. The rest of the parameters (associated with the open-loop network) as well as the type of the disturbance (including the time of entry) remain the same as in **A**. Although the output steady states change due to the presence of the disturbance, 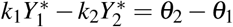 or 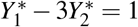 always holds (Equation (7)).

**Figure 5:**
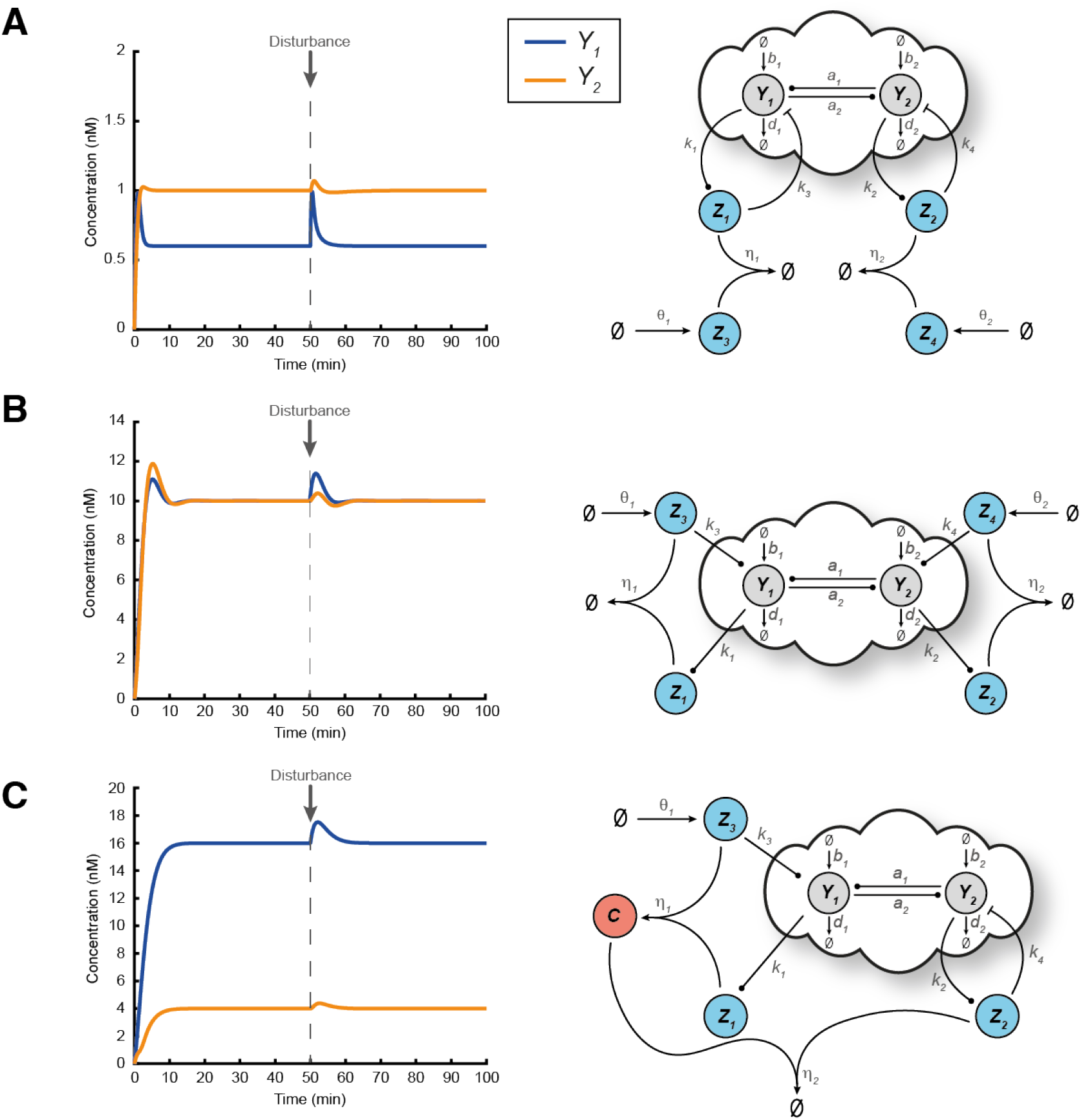
Regulating each output independently. **A** A closed-loop architecture based on the open-loop network shown in Figure 3A and D-Regulator I. For the simulated response presented here the following parameters are used: *k*_1_ = 2.5 min^−1^, *k*_2_ = 0.5 min^−1^, *k*_3_ = 2 nM^−1^ min^−1^, *k*_4_ = 2 nM^−1^ min^−1^, *η*_1_ = *η*_2_ = 10 nM^−1^ min^−1^, *θ*_1_ = 1.5 nM min^−1^, *θ*_2_ = 0.5 nM min^−1^ while the rest of the parameters (associated with the open-loop network) are the same as the ones used in Figure 3B. Despite the presence of a disturbance, 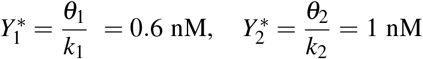 always hold (Equation (11)). **B** A closed-loop architecture based on the open-loop network shown in Figure 3A and D-Regulator II. For the simulated response presented here the following parameters are used: *k*_1_ = 1 min^−1^, *k*_2_ = 0.8 min^−1^, *k*_3_ = *k*_4_ = 0.5 min^−1^, *η*_1_ = *η*_2_ = 0.5 nM^−1^ min^−1^, *θ*_1_ = 10 nM min^−1^, *θ*_2_ = 8 nM min^−1^ while the rest of the parameters (associated with the open-loop network) are the same as the ones used in Figure 3B. Despite the presence of a disturbance, 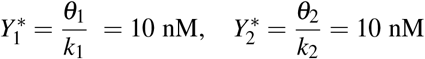 always hold (Equation (11)). **C** A closed-loop architecture based on the open-loop network shown in Figure 3A and D-Regulator III. For the simulated response presented here the following parameters are used: *k*_1_ = 0.5 min^−1^, *k*_2_ = 2 min^−1^, *k*_3_ = 0.5 min^−1^, *k*_4_ = 2 nM^−1^ min^−1^, *η*_1_ = 0.5 nM^−1^ min^−1^, *η*_2_ = 10 nM^−1^ min^−1^, *θ*_1_ = 8 nM min^−1^ while the rest of the parameters (associated with the open-loop network) are the same as the ones used in Figure 3B. Despite the presence of a disturbance, 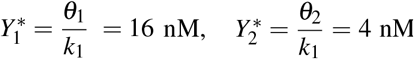 always hold (Equation (15)). The choice of the set points in **A, B** and **C** is arbitrary while the type of the disturbance (including the time of entry) is the same as in Figure 3B.

Finally, in the topology shown in Figure 5B there are two actuation reactions realized though *Z*_3_ and *Z*_4_. Due to the existence of coupling interactions in the network that we aim to control, it is evident that these actuating species act on both *Y*_1_ and *Y*_2_ simultaneously. Consequently, one could argue that an alternative way of closing the loop would be through a different species pairing (Figure 6). In particular, an annihilation (comparison) reaction between *Z*_1_, *Z*_4_ and *Z*_2_, *Z*_3_ could be used instead (*Z*_1_, *Z*_2_ can be considered as sensing species measuring the outputs *Y*_1_, *Y*_2_, respectively). However, it can be demonstrated (see Section S4 of the supplementary material) that this control strategy is not feasible since there is no realistic parameter set that can ensure closed-loop stability.

**Figure 6:**
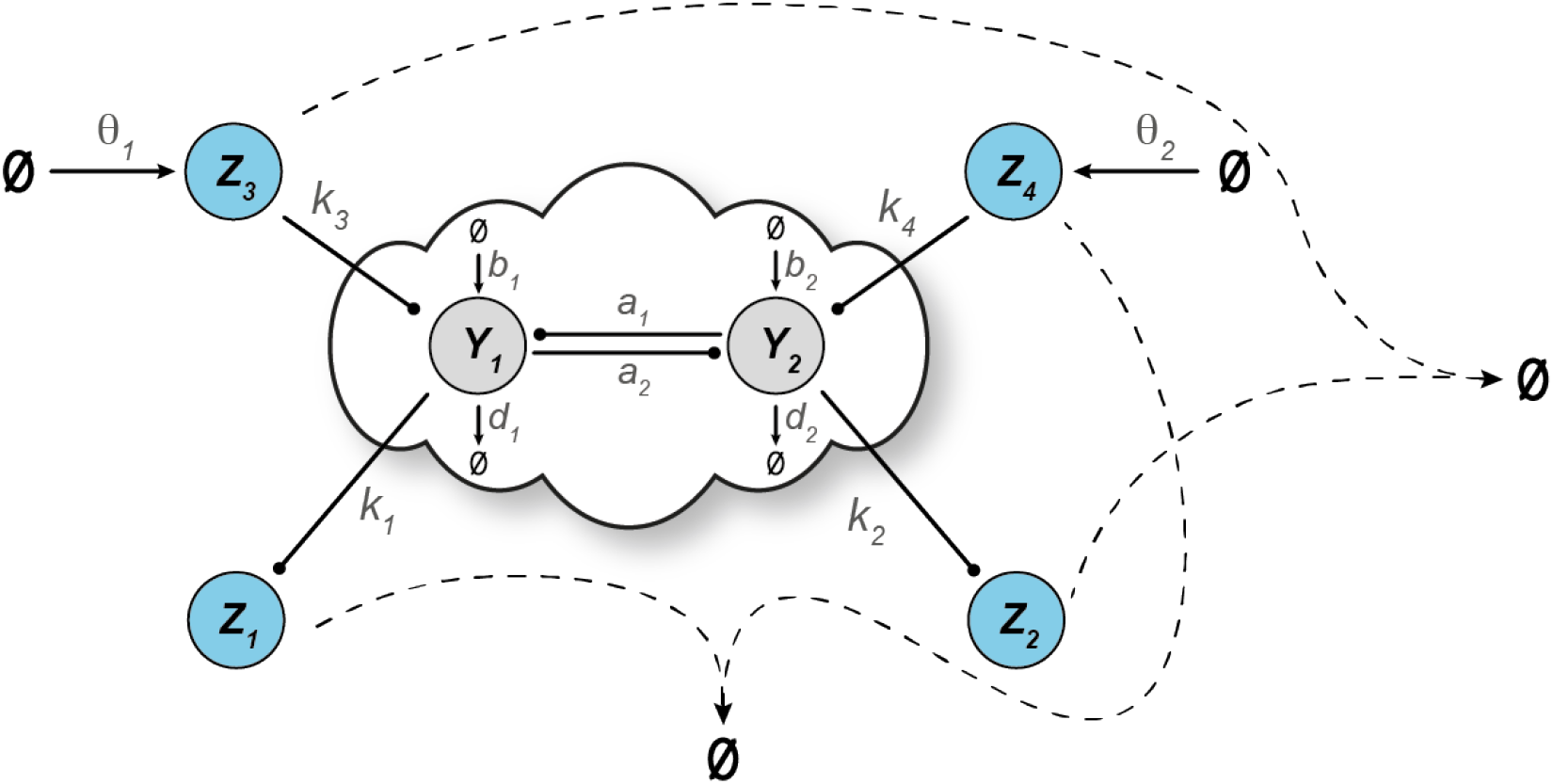
A different feedback configuration regarding the topology shown in Figure 5B which leads to instability.

### 6 Closed-loop stability

As already emphasized, assuming the existence of a finite, positive equilibrium, the proposed regulation strategies require asymptotic closed-loop stability, at least around that equilibrium (locally). A commonly used approach to assess local stability of a nonlinear system is through (Jacobian) linearization. Specifically, we can study the resulting Jacobian matrix [8]. If its eigenvalues have strictly negative real parts, i.e. the matrix is Hurwitz, then the aforementioned equilibrium is locally asymptotically stable. Necessary and sufficient conditions for that can be determined via the Routh-Hurwitz criterion (see the sections of the supplementary material associated with **Specifying the biological network to be controlled** and **Implementing the proposed regulation strategies**).

Instead of analyzing the system as a whole, we can alternatively examine it as an interconnection of two (or more) subsystems. It is often possible to assess the overall stability by studying those subsystems separately. This could be beneficial when only an input-output property of the system to be controlled is known. To demonstrate this, we consider R-Regulator and LC-Regulator with two inhibitory reactions controlling a general “cloud” network in a negative feedback configuration, as shown in Figure 1E. Focusing on the behaviour around an equilibrium of interest, we can show in both cases that if 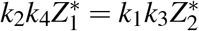 then the “controller block” corresponds to a positive real (PR) system. It is also known [33] that the negative feedback interconnection of a PR block and a weakly strictly PR (WSPR) one yields an overall asymptotically stable system. Consequently, for every WSPR “cloud block”, asymptotic closed-loop stability can be guaranteed. Further details including definitions of PR and WSPR concepts as well as proofs can be found in Section S5 of the supplementary material.

### 7 Experimental realization

To highlight the feasibility of experimentally realizing the proposed control schemes, this section describes both *in vivo* and *in vitro* implementations of the open-loop and closed-loop circuits introduced earlier. We first focus on implementations using biological parts that have been characterized in *Escherichia coli* and then discuss a molecular programming approach.

Following the description in **Specifying the biological network to be controlled**, the biological network to be controlled can be realized as shown in Figure 7. In this implementation, *Y*_1_ and *Y*_2_ are heterologous sigma factors [34], which are fused to fluorescent proteins (GFP and mCherry) to facilitate tracking of the output. Through a suitable choice of promoters, *Y*_1_ mediates the expression of *Y*_2_ and *vice versa*. Low levels of *Y*_1_ and *Y*_2_ are continuously produced from constitutive promoters, such as promoters from the BioBrick collection [35]. In all following figures, the biological parts underlying these interactions are not explicitly shown.

**Figure 7:**
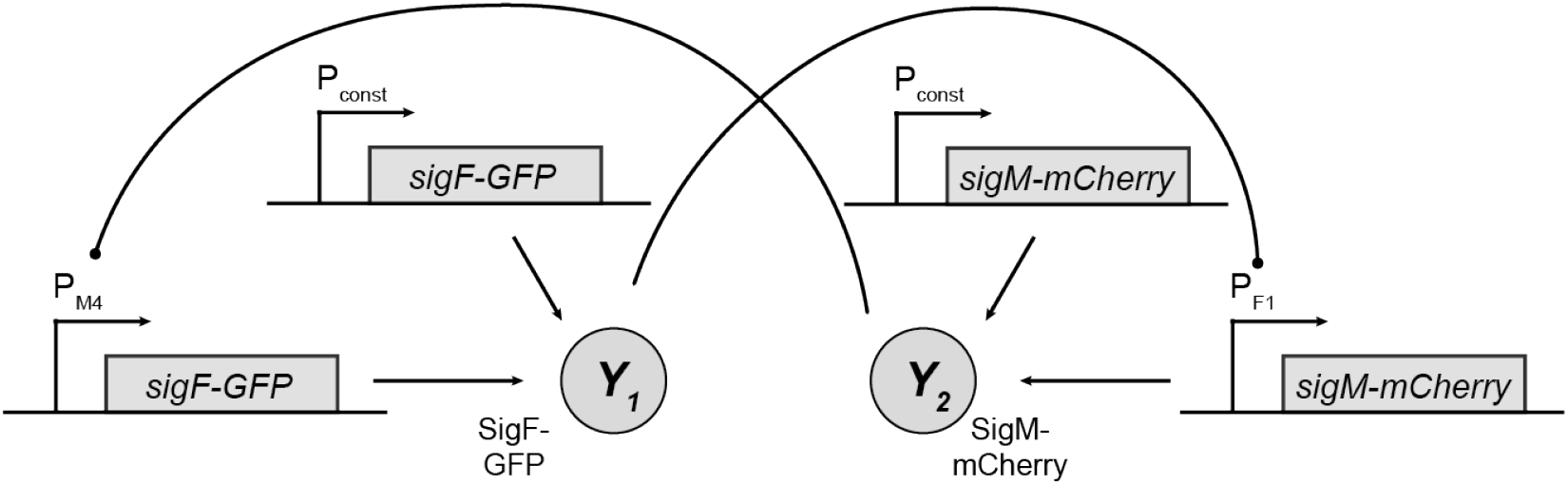
Experimental realization of the network to be controlled described by CRN (16).

#### 7.1 R-Regulator and LC-Regulator

For the proposed implementation of the R-Regulator (Figure 8), *Y*_2_ mediates expression of the hepatitis C virus protease NS3 fused to maltose-binding protein (MBP) (*Z*_2_). *Y*_1_ facilitates expression of a MBP-single-chain antibody (scFv) fusion (*Z*_1_) that specifically binds to and thus inhibits NS3 protease. Inhibition of NS3 protease activity through coexpression with single-chain antibodies in the cytoplasm of *E. coli* has been demonstrated previously [36]. Adding a suitable recognition sequence to *Y*_2_ will further allow for its degradation by NS3. An additional requirement for the LC-Regulator would be constitutive expression of *malE-scFv* and *malE-scNS3* as indicated in the dashed boxes in Figure 8.

**Figure 8:**
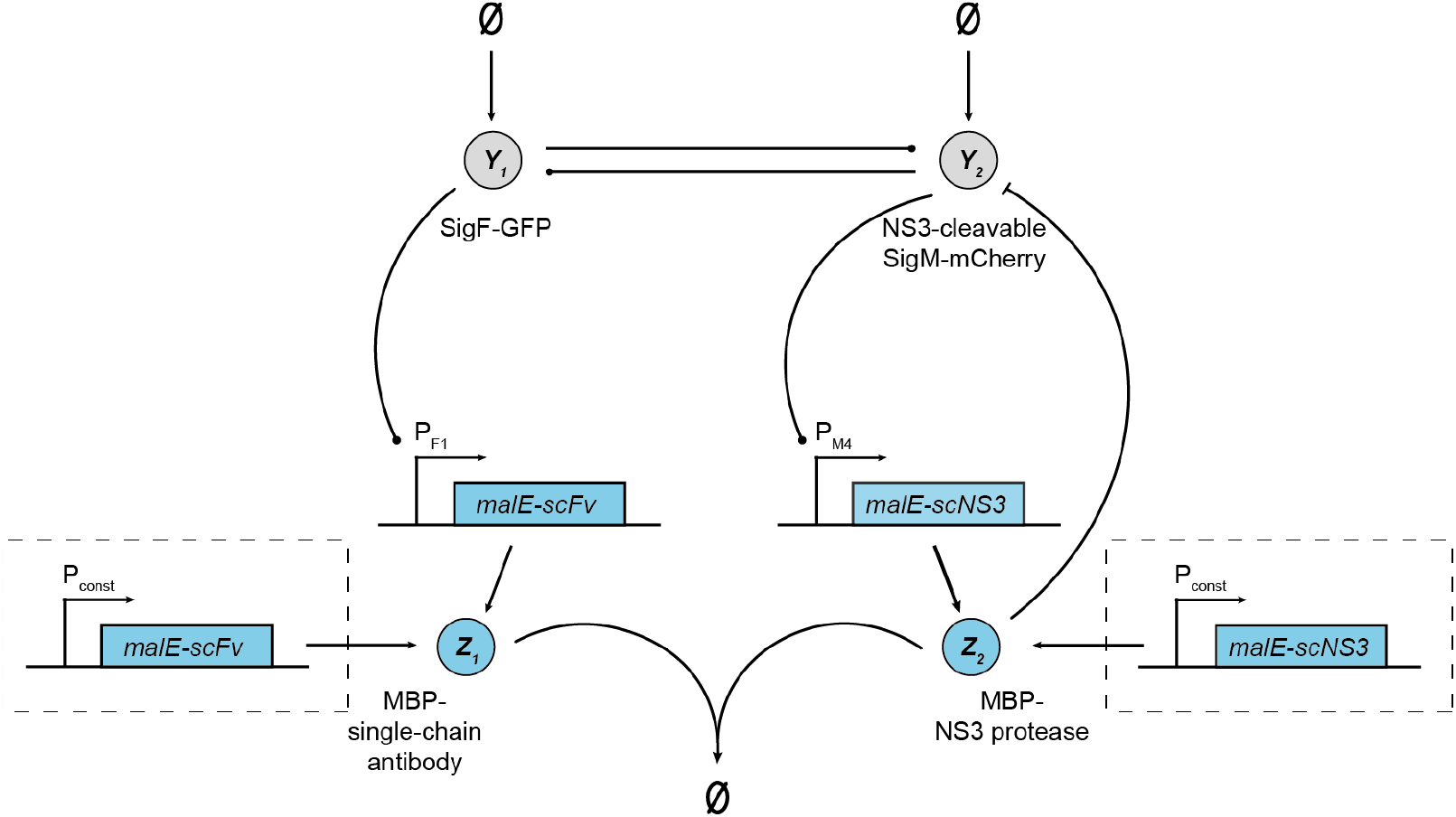
Experimental realization of the closed-loop architecture based on the open-loop network shown in Figure 6 and R-Regulator or LC-Regulator. The biological parts enclosed in dashed boxes are only required for LC-Regulator.

#### 7.2 D-Regulators

Similar to R- and LC-Regulator, the implementation for D-Regulator I makes use of the interaction between NS3 protease and a suitable single-chain antibody (Figure 9A). However, the antibody is solely expressed from a constitutive promoter in this case. As a second protease-protease inhibitor pair, we suggest use of the *E. coli* Lon protease and the phage T4 protease inhibitor PinA as discussed in our previous work [37]. For this purpose, a suitable degradation tag should be added to *Y*_1_.

**Figure 9:**
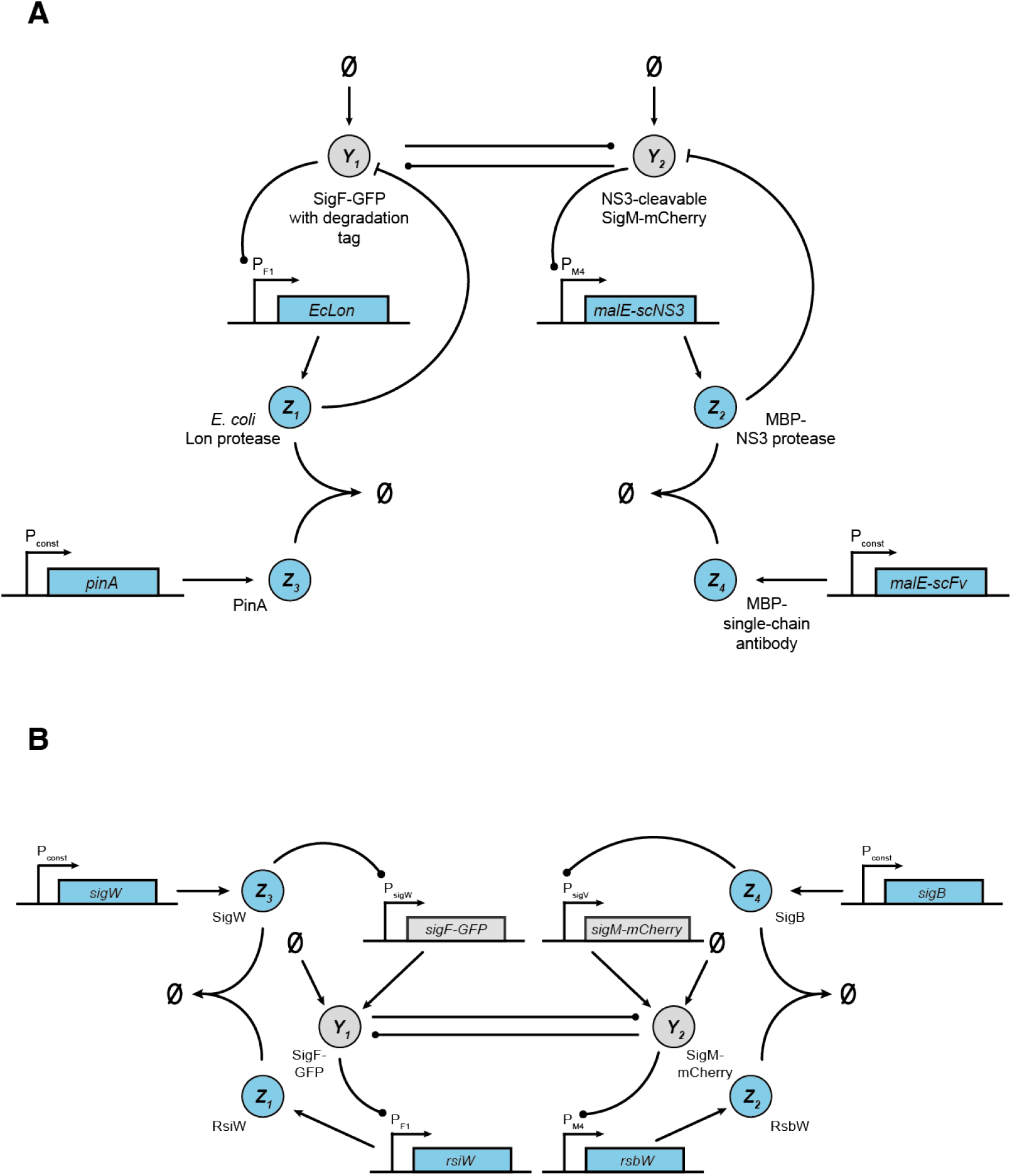
Experimental realization of the closed-loop architecture based on the open-loop network shown in Figure 6 and A D-Regulator I, B D-Regulator II.

To realize the two annihilation reactions in D-Regulator II (Figure 9B), we propose the use of *σ* - factors and anti-*σ* -factors as described previously [38, 39]. Specifically, *Z*_3_ could be the *σ* -factor SigW, which is constitutively expressed and mediates expression of SigF (*Y*_1_). SigF mediates expression on the anti-*σ* -factor RsiW (*Z*_1_), which binds to SigW. Analogous reactions are realized using SigM (*Y*_2_), SigB (*Z*_4_) and RsbW (*Z*_2_).

The design for D-Regulator III may be more difficult to implement experimentally due to the requirement of a two-stage complex formation by three biomolecules (*Z*_1_, *Z*_2_ and *Z*_3_) in addition to the requirement of *Z*_3_ catalysing the production of *Y*_1_ and *Z*_2_ inhibiting *Y*_2_. While it may be possible to achieve the desired behaviour of biomolecules using protein fusions and/or protein engineering, an alternative method to implement this design (as well as all the others) would be via molecular programming as discussed in the following section.

#### 7.3 Molecular programming implementation

In molecular programming, an abstract reaction network is realized by designing a concrete chemical reaction network using engineered molecules, so that the latter network emulates the kinetics of the former. At the edges of the abstract network, appropriate chemical transducers must be introduced to interface the abstract network with the environment. While such transducers are specific to each application, the core network is generic, and DNA (natural or synthetic) is commonly used to construct it. These systems are typically tested *in vitro* in controlled environments, with the eventual aim of embedding them in living cells, in synthetic cells [40], or in other deployable physical media. We refer to [18], Section IV, for details of concrete synthetic DNA schemes in the context of biochemical regulation, and for literature overview. Suffices to say that all the reactions used in this paper can be systematically compiled into networks of synthetic molecules that well approximate the required mass action kinetics [41].

## 8 Discussion

In this paper, we address the challenge of regulating biomolecular processes with two outputs of interest which are, in the general case, co-dependent due to coupling interactions. This co-dependence means that disturbances applied to one of the outputs will also affect the other - each of the output species may be part of a separate, independent network and, by extension, be subject to different perturbations. Thus, we propose control schemes for efficient and robust manipulation of such processes adopting concepts based on both output steady state coupling and decoupling. The proposed regulators describe biomolecular configurations with appropriate feedback interconnections which, under some assumptions, result in closed-loop systems where different types of output regulation can be achieved.

In particular, we present bio-controllers for regulating the ratio and a linear combination of the outputs referred to as R-Regulator and LC-Regulator, respectively, and three bio-controllers for regulating each of the outputs independently, namely D-Regulators I, II, III. At the core of their functioning lies a “hidden” integral feedback action realized in suitable ways in order to meet the control objectives for each case. Integral control is one of the most widely used strategies in traditional control engineering since it guarantees zero control error and constant disturbance rejection at the steady state. This comes from the fact that with this type of control, the existence of a positive/negative error, regardless of its magnitude, always generates an increasing/decreasing control signal. Essential structural components of these designs are production-inhibition loops [37] and/or annihilation reactions [29]. Moreover, to get a more practical insight, we consider a two-output biomolecular network with positive feedback coupling interactions. Treating the network as an open-loop system, we use our control designs to successfully manipulate its outputs in the presence of constant parameter perturbations. At the same time, we discuss an alternative way of closing the loop in D-Regulator-II via a different controller species “pairing”. Although it may seem reasonable, we show that this feedback configuration leads to an unstable closed-loop system.

Assuming a biologically meaningful equilibrium, the proposed designs can be used to regulate arbitrary biological processes provided that the closed-loop topologies are asymptotically stable. We therefore anticipate that they will be useful for building complex pathways that robustly respond to environmental perturbations in synthetic biology applications. To this end, we extensively discuss ways of achieving local closed-loop asymptotic stability while, for R- and LC-Regulator, we also present specific sufficient conditions based on the concept of positive realness. Furthermore, we describe possible experimental implementations of all regulators using either biomolecular species in *E. coli* or molecular programming.

Biological networks are inherently stochastic due to the probabilistic nature of biomolecular interactions [8, 42–44]. In the present study, we use deterministic mathematical analysis and simulations which offer a good approximation of the CRN dynamics when the biomolecular counts are high. Thus, an interesting future endeavour would be to investigate the behaviour of our topologies within a stochastic mathematical framework examining, for instance, both the stationary mean and variance [45–48]. Implementation of our regulatory architectures in living cells may involve an additional challenge: a decay mechanism related to cell growth, known as dilution [8], (among other factors) needs to be accounted for since it can affect the species concentrations of the controllers. Future work will therefore focus on quantifying this impact in terms of, for example, the steady state error, and explore ways to minimize it [49]. Finally, another interesting extension of our work would be to make use of dissipativity theory tools [33, 50] allowing us to study our topologies under the lens of global asymptotic stability.

## Data availability

The programming codes supporting this work can be found at: https://github.com/emgalox/MIMO-bio-controllers.

## Author contributions

Conceptualization and methodology, E.A., C.C.M.S., A.P., L.C.; Formal analysis and Software: E.A., Writing, E.A., C.C.M.S., A.P., L.C.; Supervision: A.P., L.C.

