## Supplementary Text for "Regulation strategies for two-output biomolecular networks"

Supplementary Material

Emmanouil Alexis[1], Carolin CM Schulte[1,2,#], Luca Cardelli[3], and Antonis Papachristodoulou[1,*]

[1]Department of Engineering Science, University of Oxford, Oxford OX1 3PJ, UK

[2]Department of Plant Sciences, University of Oxford, Oxford OX1 3RB, UK

[3]Department of Computer Science, University of Oxford, Oxford OX1 3QD, UK

[#]Current affiliation: Department of Biostatistics, Harvard T. H. Chan School of Public Health, Boston, MA, USA

### S1 Modelling Assumptions

The molecular interactions of the topologies presented in this work are described by chemical reaction networks (CRNs) under mass-action kinetics [1], unless otherwise stated.

### S2 Open-loop biological network

The open-loop biological network introduced in Section **4 Specifying the biological network to be controlled** of the main text is represented by the CRN (see Figure 3A) :

$$\begin{gathered}\varnothing \xrightarrow{b_1} Y_1\,, \quad \varnothing \xrightarrow{b_2} Y_2\,, \quad Y_1 \xrightarrow{d_1} \varnothing\,, \quad Y_2 \xrightarrow{d_2} \varnothing\,, \\ Y_1 \xrightarrow{\alpha_2} Y_1 + Y_2\,, \quad Y_2 \xrightarrow{\alpha_1} Y_1 + Y_2\end{gathered} \tag{S1}$$

where $b_1, b_2, d_1, d_2, \alpha_1, \alpha_2 \in \mathbb{R}_+$.

The dynamics of CRN (S1) are described by the system of Ordinary Differential Equations (ODEs):

$$\begin{bmatrix}\dot{Y}_1 \\ \dot{Y}_2\end{bmatrix} = \begin{bmatrix}-d_1 & \alpha_1 \\ \alpha_2 & -d_2\end{bmatrix}\begin{bmatrix}Y_1 \\ Y_2\end{bmatrix} + \begin{bmatrix}b_1 \\ b_2\end{bmatrix} \tag{S2}$$

Using the linear transformations $y_1 = Y_1 - Y_1^*$, $y_2 = Y_2 - Y_2^*$, we get the following mathematically equivalent system:

$$\begin{bmatrix}\dot{y}_1 \\ \dot{y}_2\end{bmatrix} = \underbrace{\begin{bmatrix}-d_1 & \alpha_1 \\ \alpha_2 & -d_2\end{bmatrix}}_{G_1}\begin{bmatrix}y_1 \\ y_2\end{bmatrix} \tag{S3}$$

where $\left(Y_1^* = \dfrac{\alpha_1 b_2 + b_1 d_2}{d_1 d_2 - \alpha_1\alpha_2}, Y_2^* = \dfrac{\alpha_2 b_1 + b_2 d_1}{d_1 d_2 - \alpha_1\alpha_2}\right)$ is the unique positive steady state of $(Y_1, Y_2)$ for any $d_1 d_2 > \alpha_1\alpha_2$.

The characteristic polynomial of system matrix $G_1$ is:

$$P_o(s) = \det(G_1 - sI) = s^2 + (d_1 + d_2)s + d_1 d_2 - \alpha_1\alpha_2$$

Both $d_1 + d_2$ and $d_1 d_2 - \alpha_1\alpha_2$ are positive and, thus, matrix $G_1$ is Hurwitz (Routh-Hurwitz criterion). Consequently, since system (S3) is linear, the origin is a globally exponentially stable steady state of (S3). By extension, $(Y_1^*, Y_2^*)$ is a globally exponentially stable steady state of system (S2).

### S3 Closed-loop biological networks

Here we analyze the behaviour of the closed-loop systems presented in Section **5 Implementing the proposed regulation strategies** of the main text.

##### R-Regulator

We have the CRN (see Figure 4A):

$$\varnothing \xrightarrow{b_1} Y_1\,,\quad \varnothing \xrightarrow{b_2} Y_2\,,\quad Y_1 \xrightarrow{d_1} \varnothing\,,\quad Y_2 \xrightarrow{d_2} \varnothing$$

$$Y_1 \xrightarrow{\alpha_2} Y_1 + Y_2\,,\quad Y_2 \xrightarrow{\alpha_1} Y_1 + Y_2\,,\quad Y_1 \xrightarrow{k_1} Y_1 + Z_1$$

$$Y_2 \xrightarrow{k_2} Y_2 + Z_2\,,\quad Y_2 + Z_2 \xrightarrow{k_3} Z_2\,,\quad Z_1 + Z_2 \xrightarrow{\eta} \varnothing$$

where $b_1$, $b_2$, $d_1$, $d_2$, $\alpha_1$, $\alpha_2$, $k_1$, $k_2$, $k_3$, $\eta \in \mathbb{R}_+$.

The corresponding ODE model is :

$$\dot{Y}_1 = b_1 - d_1 Y_1 + \alpha_1 Y_2 \qquad \text{(S4a)}$$

$$\dot{Y}_2 = b_2 - d_2 Y_2 + \alpha_2 Y_1 - k_3 Y_2 Z_2 \qquad \text{(S4b)}$$

$$\dot{Z}_1 = k_1 Y_1 - \eta Z_1 Z_2 \qquad \text{(S4c)}$$

$$\dot{Z}_2 = k_2 Y_2 - \eta Z_1 Z_2 \qquad \text{(S4d)}$$

For any $\lambda_1 = d_1 k_2 - \alpha_1 k_1 > 0$ and $\lambda_2 = b_1(\alpha_2 k_2 - d_2 k_1) + b_2(d_1 k_2 - \alpha_1 k_1) > 0$, system (S4) has a unique positive steady state:

$$Y_1^* = \frac{b_1 k_2}{\lambda_1} \qquad \text{(S5a)}$$

$$Y_2^* = \frac{b_1 k_1}{\lambda_1} \qquad \text{(S5b)}$$

$$Z_1^* = \frac{b_1^2 k_1^2 k_2 k_3}{\eta \lambda_1 \lambda_2} \qquad \text{(S5c)}$$

$$Z_2^* = \frac{\lambda_2}{b_1 k_1 k_3} \qquad \text{(S5d)}$$

Combining Equations (S5a), (S5b) yields:

$$\frac{Y_1^*}{Y_2^*} = \frac{k_2}{k_1}$$

By linearizing system (S4) around its steady state (S5) we get:

$$\begin{bmatrix} \dot{Y}_1 \\ \dot{Y}_2 \\ \dot{Z}_1 \\ \dot{Z}_2 \end{bmatrix} = \underbrace{\begin{bmatrix} -d_1 & \alpha_1 & 0 & 0 \\ \alpha_2 & -(d_2+k_3Z_2^*) & 0 & -k_3Y_2^* \\ k_1 & 0 & -\eta Z_2^* & -\eta Z_1^* \\ 0 & k_2 & -\eta Z_2^* & -\eta Z_1^* \end{bmatrix}}_{G_R} \begin{bmatrix} Y_1 \\ Y_2 \\ Z_1 \\ Z_2 \end{bmatrix} \tag{S6}$$

If all the eigenvalues of system (S6) have a negative real part - matrix $G_R$ is Hurwitz - , then (S5) is a locally exponentially stable steady state for system (S4). This stability criterion can be easily checked for a given set of parameters and was taken into account in all the simulations depicted in Section **5 Implementing the proposed regulation strategies** of the main text. Of course, as shown in this section, different closed-loop networks may result in different stability matrices. Finally, parameter regimes that guarantee local stability in each case can be found by applying the Routh-Hurwitz criterion.

#### LC-Regulator

We have the CRN (see Figure 4B):

$$\varnothing \xrightarrow{b_1} Y_1\,,\quad \varnothing \xrightarrow{b_2} Y_2\,,\quad Y_1 \xrightarrow{d_1} \varnothing\,,\quad Y_2 \xrightarrow{d_2} \varnothing \quad Y_1 \xrightarrow{\alpha_2} Y_1+Y_2$$
$$Y_2 \xrightarrow{\alpha_1} Y_1+Y_2\,,\quad \varnothing \xrightarrow{\theta_1} Z_1\,,\quad \varnothing \xrightarrow{\theta_2} Z_2 \quad Y_1 \xrightarrow{k_1} Y_1+Z_1$$
$$Y_2 \xrightarrow{k_2} Y_2+Z_2\,,\quad Y_2+Z_2 \xrightarrow{k_3} Z_2\,,\quad Z_1+Z_2 \xrightarrow{\eta} \varnothing$$

where $b_1, b_2, d_1, d_2, \alpha_1, \alpha_2, \theta_1, \theta_2, k_1, k_2, k_3, \eta \in \mathbb{R}_+$.

The corresponding ODE model is :

$$\dot{Y}_1 = b_1 - d_1Y_1 + \alpha_1Y_2 \tag{S7a}$$
$$\dot{Y}_2 = b_2 - d_2Y_2 + \alpha_2Y_1 - k_3Y_2Z_2 \tag{S7b}$$
$$\dot{Z}_1 = \theta_1 + k_1Y_1 - \eta Z_1Z_2 \tag{S7c}$$
$$\dot{Z}_2 = \theta_2 + k_2Y_2 - \eta Z_1Z_2 \tag{S7d}$$

with the following unique steady state:

$$
\begin{aligned}
Y_1^* &= \frac{\lambda_3}{\lambda_1} && \text{(S8a)}\\
Y_2^* &= \frac{\lambda_4}{\lambda_1} && \text{(S8b)}\\
Z_1^* &= \frac{k_3\lambda_4\lambda_5}{\eta\lambda_1\lambda_6} && \text{(S8c)}\\
Z_2^* &= \frac{\lambda_6}{k_3\lambda_4} && \text{(S8d)}
\end{aligned}
$$

where $\lambda_3 = b_1k_2 - \alpha_1(\theta_2-\theta_1)$, $\lambda_4 = b_1k_1 - d_1(\theta_2-\theta_1)$, $\lambda_5 = b_1k_1k_2 - \alpha_1k_1\theta_2 + d_1k_2\theta_1$, $\lambda_6 = \lambda_2 + (d_1d_2 - \alpha_1\alpha_2)(\theta_2-\theta_1)$.Here, we are interested in parameter regimes for which the steady state (S5) is positive.

Using Equations (S8a), (S8b) we calculate:

$$
k_1Y_1^* - k_2Y_2^* = \frac{1}{\lambda_1}(k_1\lambda_3 - k_2\lambda_4) \qquad \text{(S9)}
$$

Taking into account the definitions of $\lambda_1$, $\lambda_3$ and $\lambda_4$ above, relationship (S9) can be rewritten as:

$$
k_1Y_1^* - k_2Y_2^* = \frac{k_1k_2b_1 - k_1k_2b_1 + (\theta_2-\theta_1)(d_1k_2 - \alpha_1k_1)}{d_1k_2 - \alpha_1k_1}
$$

or

$$
k_1Y_1^* - k_2Y_2^* = \theta_2 - \theta_1
$$

Moreover, linearizing system (S7) around its steady state (S8) results in system (S6).

#### D-Regulator-I

We have the CRN (see Figure 5A):

$$
\varnothing \xrightarrow{b_1} Y_1\,,\quad \varnothing \xrightarrow{b_2} Y_2\,,\quad Y_1 \xrightarrow{d_1} \varnothing\,,\quad Y_2 \xrightarrow{d_2} \varnothing \quad Y_1 \xrightarrow{\alpha_2} Y_1 + Y_2
$$
$$
Y_2 \xrightarrow{\alpha_1} Y_1 + Y_2\,,\quad Y_1 \xrightarrow{k_1} Y_1 + Z_1\,,\quad Y_2 \xrightarrow{k_2} Y_2 + Z_2\,,\quad Y_1 + Z_1 \xrightarrow{k_3} Z_1
$$
$$
Y_2 + Z_2 \xrightarrow{k_4} Z_2 \quad \varnothing \xrightarrow{\theta_1} Z_3\,,\quad \varnothing \xrightarrow{\theta_2} Z_4\,,\quad Z_1 + Z_3 \xrightarrow{\eta_1} \varnothing
$$
$$
Z_2 + Z_4 \xrightarrow{\eta_2} \varnothing
$$

where $b_1, b_2, d_1, d_2, \alpha_1, \alpha_2, \theta_1, \theta_2, k_1, k_2, k_3, k_4, \eta_1, \eta_2 \in \mathbb{R}_+$.

The corresponding ODE model is :

$$
\begin{aligned}
\dot{Y}_1 &= b_1 - d_1 Y_1 + \alpha_1 Y_2 - k_3 Y_1 Z_1 && \text{(S10a)}\\
\dot{Y}_2 &= b_2 - d_2 Y_2 + \alpha_2 Y_1 - k_4 Y_2 Z_2 && \text{(S10b)}\\
\dot{Z}_1 &= k_1 Y_1 - \eta_1 Z_1 Z_3 && \text{(S10c)}\\
\dot{Z}_2 &= k_2 Y_2 - \eta_2 Z_2 Z_4 && \text{(S10d)}\\
\dot{Z}_3 &= \theta_1 - \eta_1 Z_1 Z_3 && \text{(S10e)}\\
\dot{Z}_4 &= \theta_2 - \eta_2 Z_2 Z_4 && \text{(S10f)}
\end{aligned}
$$

For any $\lambda_7 = b_1 k_1 k_2 + \alpha_1 k_1 \theta_2 - d_1 k_2 \theta_1 > 0$ and $\lambda_8 = b_2 k_1 k_2 + \alpha_2 k_2 \theta_1 - d_2 k_1 \theta_2 > 0$, system (S10) has a unique positive steady state:

$$
\begin{aligned}
Y_1^* &= \frac{\theta_1}{k_1} && \text{(S11a)}\\
Y_2^* &= \frac{\theta_2}{k_2} && \text{(S11b)}\\
Z_1^* &= \frac{\lambda_7}{k_2 k_3 \theta_1} && \text{(S11c)}\\
Z_2^* &= \frac{\lambda_8}{k_1 k_4 \theta_2} && \text{(S11d)}\\
Z_3^* &= \frac{k_2 k_3 \theta_1^2}{\eta_1 \lambda_7} && \text{(S11e)}\\
Z_4^* &= \frac{k_1 k_4 \theta_2^2}{\eta_2 \lambda_8} && \text{(S11f)}
\end{aligned}
$$

Linearization of system (S10) around its steady state (S11) gives:

$$
\begin{bmatrix} \dot{Y}_1 \\ \dot{Y}_2 \\ \dot{Z}_1 \\ \dot{Z}_2 \\ \dot{Z}_3 \\ \dot{Z}_4 \end{bmatrix}
=
\underbrace{\begin{bmatrix}
-(d_1 + k_3 Z_1^*) & \alpha_1 & -k_3 Y_1^* & 0 & 0 & 0 \\
\alpha_2 & -(d_2 + k_4 Z_2^*) & 0 & -k_4 Y_2^* & 0 & 0 \\
k_1 & 0 & -\eta_1 Z_3^* & 0 & -\eta_1 Z_1^* & 0 \\
0 & k_2 & 0 & -\eta_2 Z_4^* & 0 & -n_2 Z_2^* \\
0 & 0 & -\eta_1 Z_3^* & 0 & -\eta_1 Z_1^* & 0 \\
0 & 0 & 0 & -\eta_2 Z_4^* & 0 & -n_2 Z_2^*
\end{bmatrix}}_{G_{DI}}
\begin{bmatrix} Y_1 \\ Y_2 \\ Z_1 \\ Z_2 \\ Z_3 \\ Z_4 \end{bmatrix}
$$

#### D-Regulator-II

We have the CRN (see Figure 5B):

$$\varnothing \xrightarrow{b_1} Y_1\,, \quad \varnothing \xrightarrow{b_2} Y_2\,, \quad Y_1 \xrightarrow{d_1} \varnothing\,, \quad Y_2 \xrightarrow{d_2} \varnothing \quad Y_1 \xrightarrow{\alpha_2} Y_1+Y_2$$
$$Y_2 \xrightarrow{\alpha_1} Y_1+Y_2\,, \quad Y_1 \xrightarrow{k_1} Y_1+Z_1\,, \quad Y_2 \xrightarrow{k_2} Y_2+Z_2\,, \quad \varnothing \xrightarrow{\theta_1} Z_3$$
$$\varnothing \xrightarrow{\theta_2} Z_4\,, \quad Z_3 \xrightarrow{k_3} Z_3+Y_1 \quad Z_4 \xrightarrow{k_4} Z_4+Y_2 \quad Z_1+Z_3 \xrightarrow{\eta_1} \varnothing$$
$$Z_2+Z_4 \xrightarrow{\eta_2} \varnothing$$

where $b_1$, $b_2$, $d_1$, $d_2$, $\alpha_1$, $\alpha_2$, $\theta_1$, $\theta_2$, $k_1$, $k_2$, $k_3$, $k_4$, $\eta_1$, $\eta_2 \in \mathbb{R}_+$.

The corresponding ODE model is :

$$\dot{Y}_1 = b_1 - d_1Y_1 + \alpha_1Y_2 + k_3Z_3 \tag{S12a}$$
$$\dot{Y}_2 = b_2 - d_2Y_2 + \alpha_2Y_1 + k_4Z_4 \tag{S12b}$$
$$\dot{Z}_1 = k_1Y_1 - \eta_1Z_1Z_3 \tag{S12c}$$
$$\dot{Z}_2 = k_2Y_2 - \eta_2Z_2Z_4 \tag{S12d}$$
$$\dot{Z}_3 = \theta_1 - \eta_1Z_1Z_3 \tag{S12e}$$
$$\dot{Z}_4 = \theta_2 - \eta_2Z_2Z_4 \tag{S12f}$$

For any $\lambda_7 = b_1k_1k_2 + \alpha_1k_1\theta_2 - d_1k_2\theta_1 < 0$ and $\lambda_8 = b_2k_1k_2 + \alpha_2k_2\theta_1 - d_2k_1\theta_2 < 0$, system (S12) has a unique positive steady state:

$$Y_1^* = \frac{\theta_1}{k_1} \tag{S13a}$$
$$Y_2^* = \frac{\theta_2}{k_2} \tag{S13b}$$
$$Z_1^* = -\frac{k_1k_2k_3\theta_1}{\eta_1\lambda_7} \tag{S13c}$$
$$Z_2^* = -\frac{k_1k_2k_4\theta_2}{\eta_2\lambda_8} \tag{S13d}$$
$$Z_3^* = -\frac{\lambda_7}{k_1k_2k_3} \tag{S13e}$$
$$Z_4^* = -\frac{\lambda_8}{k_1k_2k_4} \tag{S13f}$$

We linearize system (S12) around its steady state (S13) to obtain:

$$\begin{bmatrix} \dot{Y}_1 \\ \dot{Y}_2 \\ \dot{Z}_1 \\ \dot{Z}_2 \\ \dot{Z}_3 \\ \dot{Z}_4 \end{bmatrix} = \underbrace{\begin{bmatrix} -d_1 & \alpha_1 & 0 & 0 & k_3 & 0 \\ \alpha_2 & -d_2 & 0 & 0 & 0 & k_4 \\ k_1 & 0 & -\eta_1 Z_3^* & 0 & -\eta_1 Z_1^* & 0 \\ 0 & k_2 & 0 & -\eta_2 Z_4^* & 0 & -n_2 Z_2^* \\ 0 & 0 & -\eta_1 Z_3^* & 0 & -\eta_1 Z_1^* & 0 \\ 0 & 0 & 0 & -\eta_2 Z_4^* & 0 & -n_2 Z_2^* \end{bmatrix}}_{G_{DII}} \begin{bmatrix} Y_1 \\ Y_2 \\ Z_1 \\ Z_2 \\ Z_3 \\ Z_4 \end{bmatrix}$$

#### D-Regulator-III

We have the CRN (see Figure 5C):

$$\varnothing \xrightarrow{b_1} Y_1, \quad \varnothing \xrightarrow{b_2} Y_2, \quad Y_1 \xrightarrow{d_1} \varnothing, \quad Y_2 \xrightarrow{d_2} \varnothing \quad Y_1 \xrightarrow{\alpha_2} Y_1 + Y_2$$

$$Y_2 \xrightarrow{\alpha_1} Y_1 + Y_2, \quad Y_1 \xrightarrow{k_1} Y_1 + Z_1, \quad Y_2 \xrightarrow{k_2} Y_2 + Z_2, \quad \varnothing \xrightarrow{\theta_1} Z_3$$

$$Z_3 \xrightarrow{k_3} Z_3 + Y_1, \quad Y_2 + Z_2 \xrightarrow{k_4} Z_2, \quad Z_1 + Z_3 \xrightarrow{\eta_1} C, \quad Z_2 + C \xrightarrow{\eta_2} \varnothing$$

where $b_1, b_2, d_1, d_2, \alpha_1, \alpha_2, \theta_1, k_1, k_2, k_3, k_4, \eta_1, \eta_2 \in \mathbb{R}_+$.

The corresponding ODE model is :

$$\dot{Y}_1 = b_1 - d_1 Y_1 + \alpha_1 Y_2 + k_3 Z_3 \qquad \text{(S14a)}$$

$$\dot{Y}_2 = b_2 - d_2 Y_2 + \alpha_2 Y_1 - k_4 Y_2 Z_2 \qquad \text{(S14b)}$$

$$\dot{Z}_1 = k_1 Y_1 - \eta_1 Z_1 Z_3 \qquad \text{(S14c)}$$

$$\dot{Z}_2 = k_2 Y_2 - \eta_2 Z_2 C \qquad \text{(S14d)}$$

$$\dot{Z}_3 = \theta_1 - \eta_1 Z_1 Z_3 \qquad \text{(S14e)}$$

$$\dot{C} = \eta_1 Z_1 Z_3 - \eta_2 Z_2 C \qquad \text{(S14f)}$$

For any $\lambda_9 = -b_1 k_1 k_2 + \theta_1 \lambda_1 > 0$ and $\lambda_{10} = b_2 k_1 k_2 + \theta_1(\alpha_2 k_2 - d_2 k_1) > 0$, system (S14) has a unique positive steady state:

$$
\begin{aligned}
Y_1^* &= \frac{\theta_1}{k_1} && \text{(S15a)}\\
Y_2^* &= \frac{\theta_1}{k_2} && \text{(S15b)}\\
Z_1^* &= \frac{k_1 k_2 k_3 \theta_1}{\eta_1 \lambda_9} && \text{(S15c)}\\
Z_2^* &= \frac{\lambda_{10}}{k_1 k_4 \theta_1} && \text{(S15d)}\\
Z_3^* &= \frac{\lambda_9}{k_1 k_2 k_3} && \text{(S15e)}\\
C^* &= \frac{k_1 k_4 \theta_1^2}{\eta_2 \lambda_{10}} && \text{(S15f)}
\end{aligned}
$$

Linearization of system (S14) around its steady state (S15) yields:

$$
\begin{bmatrix} \dot{Y}_1 \\ \dot{Y}_2 \\ \dot{Z}_1 \\ \dot{Z}_2 \\ \dot{Z}_3 \\ \dot{C} \end{bmatrix} = \underbrace{\begin{bmatrix} -d_1 & \alpha_1 & 0 & 0 & k_3 & 0 \\ \alpha_2 & -(d_2 + k_4 Z_2^*) & 0 & -k_4 Y_2^* & 0 & 0 \\ k_1 & 0 & -\eta_1 Z_3^* & 0 & -\eta_1 Z_1^* & 0 \\ 0 & k_2 & 0 & -\eta_2 C^* & 0 & -n_2 Z_2^* \\ 0 & 0 & -\eta_1 Z_3^* & 0 & -\eta_1 Z_1^* & 0 \\ 0 & 0 & \eta_1 Z_3^* & -\eta_2 C^* & \eta_1 Z_1^* & -n_2 Z_2^* \end{bmatrix}}_{G_{DIII}} \begin{bmatrix} Y_1 \\ Y_2 \\ Z_1 \\ Z_2 \\ Z_3 \\ C \end{bmatrix}
$$

### S4 D-Regulator-II: A different feedback configuration

Here we explore a different way of "closing the loop" in D-Regulator-II (see Section 3 **D-Regulator-II**). More specifically, we "pair" species $Z_1, Z_4$ and $Z_2, Z_3$ by assuming they can annihilate each other, respectively.

The resulting CRN is (see Figure 6):

$$
\varnothing \xrightarrow{b_1} Y_1\,, \quad \varnothing \xrightarrow{b_2} Y_2\,, \quad Y_1 \xrightarrow{d_1} \varnothing\,, \quad Y_2 \xrightarrow{d_2} \varnothing \quad Y_1 \xrightarrow{\alpha_2} Y_1 + Y_2
$$

$$
Y_2 \xrightarrow{\alpha_1} Y_1 + Y_2\,, \quad Y_1 \xrightarrow{k_1} Y_1 + Z_1\,, \quad Y_2 \xrightarrow{k_2} Y_2 + Z_2\,, \quad \varnothing \xrightarrow{\theta_1} Z_3
$$

$$
\varnothing \xrightarrow{\theta_2} Z_4\,, \quad Z_3 \xrightarrow{k_3} Z_3 + Y_1 \quad Z_4 \xrightarrow{k_4} Z_4 + Y_2 \quad Z_1 + Z_4 \xrightarrow{\eta_1} \varnothing
$$

$$
Z_2 + Z_3 \xrightarrow{\eta_2} \varnothing
$$

where $b_1, b_2, d_1, d_2, \alpha_1, \alpha_2, \theta_1, \theta_2, k_1, k_2, k_3, k_4, \eta_1, \eta_2 \in \mathbb{R}_+$.

The corresponding ODE model is :

$$\begin{aligned}
\dot{Y}_1 &= b_1 - d_1 Y_1 + \alpha_1 Y_2 + k_3 Z_3 && \text{(S16a)}\\
\dot{Y}_2 &= b_2 - d_2 Y_2 + \alpha_2 Y_1 + k_4 Z_4 && \text{(S16b)}\\
\dot{Z}_1 &= k_1 Y_1 - \eta_1 Z_1 Z_4 && \text{(S16c)}\\
\dot{Z}_2 &= k_2 Y_2 - \eta_2 Z_2 Z_3 && \text{(S16d)}\\
\dot{Z}_3 &= \theta_1 - \eta_2 Z_2 Z_3 && \text{(S16e)}\\
\dot{Z}_4 &= \theta_2 - \eta_1 Z_1 Z_4 && \text{(S16f)}
\end{aligned}$$

For any $\lambda_{11} = d_2 k_1 \theta_1 - \alpha_2 k_2 \theta_2 - b_2 k_1 k_2 > 0$ and $\lambda_{12} = d_1 k_2 \theta_2 - \alpha_1 k_1 \theta_1 - b_1 k_1 k_2 > 0$, system (S16) has a unique positive steady state:

$$\begin{aligned}
Y_1^* &= \frac{\theta_2}{k_1} && \text{(S17a)}\\
Y_2^* &= \frac{\theta_1}{k_2} && \text{(S17b)}\\
Z_1^* &= \frac{k_1 k_2 k_4 \theta_2}{\eta_1 \lambda_{11}} && \text{(S17c)}\\
Z_2^* &= \frac{k_1 k_2 k_3 \theta_1}{\eta_2 \lambda_{12}} && \text{(S17d)}\\
Z_3^* &= \frac{\lambda_{12}}{k_1 k_2 k_3} && \text{(S17e)}\\
Z_4^* &= \frac{\lambda_{11}}{k_1 k_2 k_4} && \text{(S17f)}
\end{aligned}$$

By linearizing system (S16) around its steady state (S17) we get:

$$\begin{bmatrix} \dot{Y}_1 \\ \dot{Y}_2 \\ \dot{Z}_1 \\ \dot{Z}_2 \\ \dot{Z}_3 \\ \dot{Z}_4 \end{bmatrix} = \underbrace{\begin{bmatrix} -d_1 & \alpha_1 & 0 & 0 & k_3 & 0 \\ \alpha_2 & -d_2 & 0 & 0 & 0 & k_4 \\ k_1 & 0 & -\mu_{14} & 0 & 0 & -\mu_{11} \\ 0 & k_2 & 0 & -\mu_{23} & -\mu_{22} & 0 \\ 0 & 0 & 0 & -\mu_{23} & -\mu_{22} & 0 \\ 0 & 0 & -\mu_{14} & 0 & 0 & -\mu_{11} \end{bmatrix}}_{G_{DFII}} \begin{bmatrix} Y_1 \\ Y_2 \\ Z_1 \\ Z_2 \\ Z_3 \\ Z_4 \end{bmatrix} \qquad \text{(S18)}$$

where $\mu_{14} = \eta_1 Z_4^*$, $\mu_{11} = \eta_1 Z_1^*$, $\mu_{23} = \eta_2 Z_3^*$ and $\mu_{22} = \eta_2 Z_2^*$.

The determinant of matrix $G_{DFII}$ can be calculated as follows:

$$\det G_{DFII} = \begin{vmatrix} -d_1 & \alpha_1 & 0 & 0 & k_3 & 0 \\ \alpha_2 & -d_2 & 0 & 0 & 0 & k_4 \\ k_1 & 0 & -\mu_{14} & 0 & 0 & -\mu_{11} \\ 0 & k_2 & 0 & -\mu_{23} & -\mu_{22} & 0 \\ 0 & 0 & 0 & -\mu_{23} & -\mu_{22} & 0 \\ 0 & 0 & -\mu_{14} & 0 & 0 & -\mu_{11} \end{vmatrix} = (-1) \begin{vmatrix} k_1 & 0 & -\mu_{14} & 0 & 0 & -\mu_{11} \\ \alpha_2 & -d_2 & 0 & 0 & 0 & k_4 \\ -d_1 & \alpha_1 & 0 & 0 & k_3 & 0 \\ 0 & k_2 & 0 & -\mu_{23} & -\mu_{22} & 0 \\ 0 & 0 & 0 & -\mu_{23} & -\mu_{22} & 0 \\ 0 & 0 & -\mu_{14} & 0 & 0 & -\mu_{11} \end{vmatrix}$$

$$= (-1) \begin{vmatrix} k_1 & 0 & -\mu_{14} & 0 & 0 & -\mu_{11} \\ 0 & -d_2 & \frac{\alpha_2 \mu_{14}}{k_1} & 0 & 0 & \frac{k_1 k_4 + \alpha_2 \mu_{11}}{k_1} \\ -d_1 & \alpha_1 & 0 & 0 & k_3 & 0 \\ 0 & k_2 & 0 & -\mu_{23} & -\mu_{22} & 0 \\ 0 & 0 & 0 & -\mu_{23} & -\mu_{22} & 0 \\ 0 & 0 & -\mu_{14} & 0 & 0 & -\mu_{11} \end{vmatrix} = (-1) \begin{vmatrix} k_1 & 0 & -\mu_{14} & 0 & 0 & -\mu_{11} \\ 0 & -d_2 & \frac{\alpha_2 \mu_{14}}{k_1} & 0 & 0 & \frac{k_1 k_4 + \alpha_2 \mu_{11}}{k_1} \\ 0 & \alpha_1 & -\frac{d_1 \mu_{14}}{k_1} & 0 & k_3 & -\frac{d_1 \mu_{11}}{k_1} \\ 0 & k_2 & 0 & -\mu_{23} & -\mu_{22} & 0 \\ 0 & 0 & 0 & -\mu_{23} & -\mu_{22} & 0 \\ 0 & 0 & -\mu_{14} & 0 & 0 & -\mu_{11} \end{vmatrix}$$

$$= (-1)^2 \begin{vmatrix} k_1 & 0 & -\mu_{14} & 0 & 0 & -\mu_{11} \\ 0 & k_2 & 0 & -\mu_{23} & -\mu_{22} & 0 \\ 0 & \alpha_1 & -\frac{d_1 \mu_{14}}{k_1} & 0 & k_3 & -\frac{d_1 \mu_{11}}{k_1} \\ 0 & -d_2 & \frac{\alpha_2 \mu_{14}}{k_1} & 0 & 0 & \frac{k_1 k_4 + \alpha_2 \mu_{11}}{k_1} \\ 0 & 0 & 0 & -\mu_{23} & -\mu_{22} & 0 \\ 0 & 0 & -\mu_{14} & 0 & 0 & -\mu_{11} \end{vmatrix} = (-1)^2 \begin{vmatrix} k_1 & 0 & -\mu_{14} & 0 & 0 & -\mu_{11} \\ 0 & k_2 & 0 & -\mu_{23} & -\mu_{22} & 0 \\ 0 & 0 & -\frac{d_1 \mu_{14}}{k_1} & \frac{\alpha_1 \mu_{23}}{k_2} & \frac{k_2 k_3 + \alpha_1 \mu_{22}}{k_2} & -\frac{d_1 \mu_{11}}{k_1} \\ 0 & -d_2 & \frac{\alpha_2 \mu_{14}}{k_1} & 0 & 0 & \frac{k_1 k_4 + \alpha_2 \mu_{11}}{k_1} \\ 0 & 0 & 0 & -\mu_{23} & -\mu_{22} & 0 \\ 0 & 0 & -\mu_{14} & 0 & 0 & -\mu_{11} \end{vmatrix}$$

$$=(-1)^2\begin{vmatrix} k_1 & 0 & -\mu_{14} & 0 & 0 & -\mu_{11} \\ 0 & k_2 & 0 & -\mu_{23} & -\mu_{22} & 0 \\ 0 & 0 & -\frac{d_1\mu_{14}}{k_1} & \frac{\alpha_1\mu_{23}}{k_2} & \frac{k_2k_3+\alpha_1\mu_{22}}{k_2} & -\frac{d_1\mu_{11}}{k_1} \\ 0 & 0 & \frac{\alpha_2\mu_{14}}{k_1} & -\frac{d_2\mu_{23}}{k_2} & -\frac{d_2\mu_{22}}{k_2} & \frac{k_1k_4+\alpha_2\mu_{11}}{k_1} \\ 0 & 0 & 0 & -\mu_{23} & -\mu_{22} & 0 \\ 0 & 0 & -\mu_{14} & 0 & 0 & -\mu_{11} \end{vmatrix}$$

$$=(-1)^3\begin{vmatrix} k_1 & 0 & -\mu_{14} & 0 & 0 & -\mu_{11} \\ 0 & k_2 & 0 & -\mu_{23} & -\mu_{22} & 0 \\ 0 & 0 & -\mu_{14} & 0 & 0 & -\mu_{11} \\ 0 & 0 & \frac{\alpha_2\mu_{14}}{k_1} & -\frac{d_2\mu_{23}}{k_2} & -\frac{d_2\mu_{22}}{k_2} & \frac{k_1k_4+\alpha_2\mu_{11}}{k_1} \\ 0 & 0 & 0 & -\mu_{23} & -\mu_{22} & 0 \\ 0 & 0 & -\frac{d_1\mu_{14}}{k_1} & \frac{\alpha_1\mu_{23}}{k_2} & \frac{k_2k_3+\alpha_1\mu_{22}}{k_2} & -\frac{d_1\mu_{11}}{k_1} \end{vmatrix}$$

$$=(-1)^3\begin{vmatrix} k_1 & 0 & -\mu_{14} & 0 & 0 & -\mu_{11} \\ 0 & k_2 & 0 & -\mu_{23} & -\mu_{22} & 0 \\ 0 & 0 & -\mu_{14} & 0 & 0 & -\mu_{11} \\ 0 & 0 & 0 & -\frac{d_2\mu_{23}}{k_2} & -\frac{d_2\mu_{22}}{k_2} & k_4 \\ 0 & 0 & 0 & -\mu_{23} & -\mu_{22} & 0 \\ 0 & 0 & -\frac{d_1\mu_{14}}{k_1} & \frac{\alpha_1\mu_{23}}{k_2} & \frac{k_2k_3+\alpha_1\mu_{22}}{k_2} & -\frac{d_1\mu_{11}}{k_1} \end{vmatrix} =(-1)^3\begin{vmatrix} k_1 & 0 & -\mu_{14} & 0 & 0 & -\mu_{11} \\ 0 & k_2 & 0 & -\mu_{23} & -\mu_{22} & 0 \\ 0 & 0 & -\mu_{14} & 0 & 0 & -\mu_{11} \\ 0 & 0 & 0 & -\frac{d_2\mu_{23}}{k_2} & -\frac{d_2\mu_{22}}{k_2} & k_4 \\ 0 & 0 & 0 & -\mu_{23} & -\mu_{22} & 0 \\ 0 & 0 & 0 & \frac{\alpha_1\mu_{23}}{k_2} & \frac{k_2k_3+\alpha_1\mu_{22}}{k_2} & 0 \end{vmatrix}$$

$$=(-1)^4\begin{vmatrix} k_1 & 0 & -\mu_{14} & 0 & 0 & -\mu_{11} \\ 0 & k_2 & 0 & -\mu_{23} & -\mu_{22} & 0 \\ 0 & 0 & -\mu_{14} & 0 & 0 & -\mu_{11} \\ 0 & 0 & 0 & \frac{\alpha_1\mu_{23}}{k_2} & \frac{k_2k_3+\alpha_1\mu_{22}}{k_2} & 0 \\ 0 & 0 & 0 & -\mu_{23} & -\mu_{22} & 0 \\ 0 & 0 & 0 & -\frac{d_2\mu_{23}}{k_2} & -\frac{d_2\mu_{22}}{k_2} & k_4 \end{vmatrix} =(-1)^4\begin{vmatrix} k_1 & 0 & -\mu_{14} & 0 & 0 & -\mu_{11} \\ 0 & k_2 & 0 & -\mu_{23} & -\mu_{22} & 0 \\ 0 & 0 & -\mu_{14} & 0 & 0 & -\mu_{11} \\ 0 & 0 & 0 & \frac{\alpha_1\mu_{23}}{k_2} & \frac{k_2k_3+\alpha_1\mu_{22}}{k_2} & 0 \\ 0 & 0 & 0 & 0 & \frac{k_2k_3}{\alpha_1} & 0 \\ 0 & 0 & 0 & -\frac{d_2\mu_{23}}{k_2} & -\frac{d_2\mu_{22}}{k_2} & k_4 \end{vmatrix}$$

$$
= (-1)^4 \begin{vmatrix} k_1 & 0 & -\mu_{14} & 0 & 0 & -\mu_{11} \\ 0 & k_2 & 0 & -\mu_{23} & -\mu_{22} & 0 \\ 0 & 0 & -\mu_{14} & 0 & 0 & -\mu_{11} \\ 0 & 0 & 0 & \frac{\alpha_1\mu_{23}}{k_2} & \frac{k_2k_3+\alpha_1\mu_{22}}{k_2} & 0 \\ 0 & 0 & 0 & 0 & \frac{k_2k_3}{\alpha_1} & 0 \\ 0 & 0 & 0 & 0 & \frac{d_2k_3}{\alpha_1} & k_4 \end{vmatrix} = (-1)^5 \begin{vmatrix} k_1 & 0 & -\mu_{14} & 0 & 0 & -\mu_{11} \\ 0 & k_2 & 0 & -\mu_{23} & -\mu_{22} & 0 \\ 0 & 0 & -\mu_{14} & 0 & 0 & -\mu_{11} \\ 0 & 0 & 0 & \frac{\alpha_1\mu_{23}}{k_2} & \frac{k_2k_3+\alpha_1\mu_{22}}{k_2} & 0 \\ 0 & 0 & 0 & 0 & \frac{d_2k_3}{\alpha_1} & k_4 \\ 0 & 0 & 0 & 0 & \frac{k_2k_3}{\alpha_1} & 0 \end{vmatrix}
$$

$$
= (-1)^5 \begin{vmatrix} k_1 & 0 & -\mu_{14} & 0 & 0 & -\mu_{11} \\ 0 & k_2 & 0 & -\mu_{23} & -\mu_{22} & 0 \\ 0 & 0 & -\mu_{14} & 0 & 0 & -\mu_{11} \\ 0 & 0 & 0 & \frac{\alpha_1\mu_{23}}{k_2} & \frac{k_2k_3+\alpha_1\mu_{22}}{k_2} & 0 \\ 0 & 0 & 0 & 0 & \frac{d_2k_3}{\alpha_1} & k_4 \\ 0 & 0 & 0 & 0 & 0 & -\frac{k_2k_4}{d_2} \end{vmatrix}
$$

or

$$
\det G_{DFII} = -k_1k_2k_3k_4\mu_{14}\mu_{23} \tag{S19}
$$

since the determinant of a triangular matrix is equal to the product of the entries on the main diagonal.

Note also the following:

- The degree of the characteristic polynomial of matrix $G_{DFII}$, $P_{DFII}(s) = \det(G_{DFII} - sI)$, and, by extension, the number of the eigenvalues of system (S18) is 6 (counting multiplicities).
- The product of the eigenvalues of system (S18) is equal to $detG_{DFII}$.
- All the entries of matrix $G_{DFII}$ are real. Consequently, its complex eigenvalues (if they exist) occur in conjugate pairs.
- The product of a complex number and its conjugate is a real, non-negative number.
- As already discussed, the parameters and the steady state of the system under consideration are positive. Thus, Equation (S19) indicates that $\det G_{DFII} < 0$

We therefore conclude that at least one of the eigenvalues of system (S18) is real and positive. This implies that steady state (S17) cannot be stable and, thus, the feedback configuration in question does not constitute an efficient regulation strategy.

### S5 Feedback interconnection and closed-loop stability

#### Useful mathematical concepts

- Here we deal with linear, time-invariant systems whose input-output relationship in the Laplace domain can be described by a proper, rational and square transfer function matrix $H(s)$, where $s \in \mathbb{C}$ is the Laplace variable.

- A state-space realization $(A, B, C, D)$ of $H(s)$ is said to be a minimal realization of $H(s)$ if $A$ has the smallest possible dimension (i.e. the fewest number of states). The smallest dimension is called the McMillan degree of $H(s)$. A mode is hidden if it is not state controllable or observable and thus does not appear in the minimal realization. (Definition 4.3 in [2]) Moreover, a state-space realization is minimal if and only if $(A, B)$ is state controllable and $(A, C)$ is state observable [2]. Here, we consider only such state-space realizations.

- The transfer matrix $H(s) \in \mathbb{C}^{m\times m}$ is positive real (PR) if i) $H(s)$ has no pole in $\mathbf{Re}[s] > 0$, ii) $H(s)$ is real for all positive real $s$, iii) $H(s) + H^H(s) \succcurlyeq 0$ for all $\mathbf{Re}[s] > 0$. (Definition 2.34 in [3]).

- A rational transfer matrix $H(s) \in \mathbb{C}^{m\times m}$ is weakly strictly positive real (WSPR) if i) $H(s)$ is analytic in $\mathbf{Re}[s] \geq 0$, ii) $H(j\omega) + H^T(-j\omega) \succ 0$ for all $\omega \in \mathbb{R}$. (Definition 2.77 in [3]).

The notations $^H$ and $^T$ indicate the conjugate transpose and the transpose of a matrix, respectively while $\succ$ ($\succcurlyeq$) indicates a positive definite (positive-semidefinite) matrix.

#### R- and LC- Regulator and closed-loop behaviour

We consider a general ("cloud") biomolecular process (see Figure 1A) consisting of $q$ species, $Y_1, Y_2, \ldots, Y_q$ which participate in an arbitrary number of chemical reactions following mass action kinetics. The dynamics of the process can be represented as:

$$\dot{Y} = f(Y) \tag{S20}$$

where $Y = [Y_1 \;\; Y_2 \;\; \dots \;\; Y_q]^T$. Species $Y_1, Y_2$ are treated as the species of interest.

We now consider the feedback configuration depicted in Figure 1E where R-Regulator and LC-Regulator are used to control the target (output) species, $Y_1, Y_2$, of the aforementioned "cloud" process. Given the analysis of Section **2 Control schemes with steady state coupling** of the main text and Equation (S20), we have for the closed-loop dynamics:

- R-Regulator case

$$\begin{aligned} \dot{Y} &= f(Y) - \xi_1 k_4 Y_1 Z_1 - \xi_2 k_3 Y_2 Z_2 && \text{(S21a)} \\ \dot{Z}_1 &= k_1 Y_1 - \eta_1 Z_1 Z_3 && \text{(S21b)} \\ \dot{Z}_2 &= k_2 Y_2 - \eta_2 Z_2 Z_4 && \text{(S21c)} \\ & && \text{(S21d)} \end{aligned}$$

- LC-Regulator case

$$\begin{aligned} \dot{Y} &= f(Y) - \xi_1 k_4 Y_1 Z_1 - \xi_2 k_3 Y_2 Z_2 && \text{(S22a)} \\ \dot{Z}_1 &= \theta_1 + k_1 Y_1 - \eta_1 Z_1 Z_3 && \text{(S22b)} \\ \dot{Z}_2 &= \theta_2 + k_2 Y_2 - \eta_2 Z_2 Z_4 && \text{(S22c)} \\ & && \text{(S22d)} \end{aligned}$$

where $\xi_1 = [1 \;\; 0 \;\; \dots \;\; 0]^T$, $\xi_2 = [0 \;\; 1 \;\; \dots \;\; 0]^T \in \mathbb{Z}^q$ and $b_1$, $b_2$, $d_1$, $d_2$, $\alpha_1$, $\alpha_2$, $\theta_1$, $\theta_2$, $k_1$, $k_2$, $k_3$, $k_4$, $\eta$ $\in \mathbb{R}_+$.

We assume a finite, positive steady state (equilibrium) of interest $E = (Y_1^*, Y_2^*, \dots, Y_q^*, Z_1^*, Z_2^*)$ and we focus on the behaviour of the above closed-loop systems around it. We therefore adopt the coordinate transformations $y_1 = Y_1 - Y_1^*, y_2 = Y_2 - Y_2^*, \dots, y_q = Y_q - Y_q^*, z_1 = Z_1 - Z_1^*, z_2 = Z_2 - Z_2^*$ which denote small perturbations around the aforementioned steady state. The resulting linearized dynamics of both systems (S21) and (S22) are described as:

$$\begin{bmatrix} \dot{y} \\ \dot{z}_1 \\ \dot{z}_2 \end{bmatrix} = \begin{bmatrix} A_p & -\xi_1 k_4 Y_1^* & -\xi_2 k_3 Y_2^* \\ \xi_1^T k_1 & -\eta Z_2^* & -\eta Z_1^* \\ \xi_2^T k_2 & -\eta Z_2^* & -\eta Z_1^* \end{bmatrix} \begin{bmatrix} y \\ z_1 \\ z_2 \end{bmatrix} \tag{S23}$$

where $y = [y_1 \;\; y_2 \;\; \ldots \;\; y_q]^T$ and $A_p = \left.\dfrac{\partial f}{\partial y}\right|_E - \xi_1 k_4 Z_1^* - \xi_2 k_3 Z_2^*$.

System (S23) can be seen as the negative feedback interconnection of two subsystems representing the (linearized) "cloud" process and the controller, respectively. More specifically, we have:

$$\dot{y} = A_p y + B_p u_p \tag{S24a}$$

$$w_p = C_p y + D_p u_p \tag{S24b}$$

and

$$\dot{z} = A_c z + B_c u_c \tag{S25a}$$

$$w_c = C_c z + D_c u_c \tag{S25b}$$

where $z = [z_1 \;\; z_2]^T$, $u_p = [u_{1p} \;\; u_{2p}]^T$, $u_c = [u_{1c} \;\; u_{2c}]^T$, $A_c = \begin{bmatrix} -\eta Z_2^* & -\eta Z_1^* \\ -\eta Z_2^* & -\eta Z_1^* \end{bmatrix}$ $B_p = [\xi_1 \;\; \xi_2]$, $B_c = \begin{bmatrix} k_1 & 0 \\ 0 & k_2 \end{bmatrix}$, $C_p = [\xi_1 \;\; \xi_2]^T$, $C_c = \begin{bmatrix} k_4 & 0 \\ 0 & k_3 \end{bmatrix}$, $D_p = D_c = 0$. In addition, $u_p = -w_c$ and $u_c = w_p$.

We now calculate the transfer function matrix corresponding to state-space model (S25) as $H_c(s) = C_c(sI - A_c)^{-1} + D_c$ to obtain:

$$H_c(s) = \begin{bmatrix} \frac{k_1 k_4 (s+\eta Z_1^*)}{s(s+\eta(Z_1^*+Z_2^*))} & \frac{-k_2 k_4 \eta Z_1^*}{s(s+\eta(Z_1^*+Z_2^*))} \\ \frac{-k_1 k_3 \eta Z_2^*}{s(s+\eta(Z_1^*+Z_2^*))} & \frac{k_2 k_3 (s+\eta Z_2^*)}{s(s+\eta(Z_1^*+Z_2^*))} \end{bmatrix} \tag{S26}$$

Here $W_c(s) = H_c(s) U_c(s)$, where $W_c(s)$ and $U_c(s)$ are the Laplace transform of $w_c$ and $u_c$, respectively.

**Theorem** If $k_2 k_4 Z_1^* = k_1 k_3 Z_2^*$, then the transfer function matrix $H_c(s)$ (Equation (S26)) is positive real (PR).

*Proof.* For $k_2 k_4 Z_1^* = k_1 k_3 Z_2^*$, Equation (S26) can be written as:

$$H_c(s) = \begin{bmatrix} \frac{k_1 k_4 (s+\eta Z_1^*)}{s(s+\eta(Z_1^*+Z_2^*))} & \frac{-k_2 k_4 \eta Z_1^*}{s(s+\eta(Z_1^*+Z_2^*))} \\ \frac{-k_2 k_4 \eta Z_1^*}{s(s+\eta(Z_1^*+Z_2^*))} & \frac{k_2 k_3 (s+\eta \frac{k_2 k_4}{k_1 k_3} Z_1^*)}{s(s+\eta(Z_1^*+Z_2^*))} \end{bmatrix} \tag{S27}$$

Transfer function matrix (S27) has no poles in $\mathbf{Re}[s] > 0$.

We also calculate:

$$H_c(j\omega) + H_c^H(j\omega) = \begin{bmatrix} \frac{1}{k_3}\frac{2k_2k_4^2\eta Z_1^*}{\omega^2+\eta^2(Z_1^*+Z_2^*)^2} & \frac{2k_2k_4\eta Z_1^*}{\omega^2+\eta^2(Z_1^*+Z_2^*)^2} \\ \frac{2k_2k_4\eta Z_1^*}{\omega^2+\eta^2(Z_1^*+Z_2^*)^2} & \frac{2k_2k_3\eta Z_1^*}{\omega^2+\eta^2(Z_1^*+Z_2^*)^2} \end{bmatrix}$$

$H_c(j\omega) + H_c^H(j\omega) \succcurlyeq 0$ since $tr\big(H_c(j\omega) + H_c^H(j\omega)\big) > 0$ and $det\big(H_c(j\omega) + H_c^H(j\omega)\big) = 0$ for all $\omega$.

In addition, $j\omega_0$ is a simple pole of transfer function matrix (S27) with $\omega_0 = 0$ while the corresponding residual is:

$$K_0 = \lim_{s\to 0} sH_c(s) = \begin{bmatrix} \frac{k_1k_4\eta Z_1^*}{\eta(Z_1^*+Z_2^*)} & \frac{-k_2k_4\eta Z_1^*}{\eta(Z_1^*+Z_2^*)} \\ \frac{-k_2k_4\eta Z_1^*}{\eta(Z_1^*+Z_2^*)} & \frac{k_2^2k_4\eta Z_1^*}{k_1\eta(Z_1^*+Z_2^*)} \end{bmatrix}$$

$K_0 \succcurlyeq 0$ since $tr(K_0) > 0$ and $det(K_0) = 0$.

Thus, according to Theorem 2.48 in [3] transfer function matrix (S27) is PR. □

Let now $H_p(s)$ be the transfer function matrix corresponding to state-space model (S24). According to Lemma 3.67 in [3], if $H_p(s)$ is WSPR, then the closed-loop system (S23) is asymptotically stable.

### Toy example

We consider a closed-loop system based on R-Regulator described by the following CRN:

$$\varnothing \xrightarrow{b_1} Y_1\,, \quad \varnothing \xrightarrow{b_2} Y_2\,, \quad Y_1 \xrightarrow{k_1} Y_1 + Z_1$$
$$Y_2 \xrightarrow{k_2} Y_2 + Z_2\,, \quad Z_1 + Z_2 \xrightarrow{\eta} \varnothing$$
$$Y_1 + Z_1 \xrightarrow{k_4} Z_1\,, \quad Y_2 + Z_2 \xrightarrow{k_3} Z_2$$

For simplicity, we assume unitary kinetic parameter values and obtain the following ODE model for the dynamics:

$$\dot{Y}_1 = 1 - Y_1Z_1 \qquad \text{(S28a)}$$

$$\dot{Y}_2 = 1 - Y_2Z_2 \qquad \text{(S28b)}$$

$$\dot{Z}_1 = Y_1 - Z_1Z_2 \qquad \text{(S28c)}$$

$$\dot{Z}_2 = Y_2 - Z_1Z_2 \qquad \text{(S28d)}$$

The point $E = (1,1,1,1)$ is a steady-state for system (S28). The linearized dynamics about $E$ is given by:

$$\begin{bmatrix} \dot{y}_1 \\ \dot{y}_2 \\ \dot{z}_1 \\ \dot{z}_2 \end{bmatrix} = \begin{bmatrix} -1 & 0 & -1 & 0 \\ 0 & -1 & 0 & -1 \\ 1 & 0 & -1 & -1 \\ 0 & 1 & -1 & -1 \end{bmatrix} \begin{bmatrix} y_1 \\ y_2 \\ z_1 \\ z_2 \end{bmatrix} \tag{S29}$$

As can be seen, $k_2 k_4 Z_1^* = k_1 k_3 Z_2^* = 1$.

Moreover:

$$H_p(s) = C_p(sI - A_p)^{-1} + D_p = \begin{bmatrix} \frac{1}{s+1} & 0 \\ 0 & \frac{1}{s+1} \end{bmatrix} \tag{S30}$$

which is analytic in $\mathbf{Re}[s] \geq 0$.

We also calculate:

$$H_p(j\omega) + H_p^H(j\omega) = \begin{bmatrix} \frac{2}{\omega^2+1} & 0 \\ 0 & \frac{2}{\omega^2+1} \end{bmatrix}$$

$H_p(j\omega) + H_p^H(j\omega) \succcurlyeq 0$ since $tr\big(H_c(j\omega) + H_c^H(j\omega)\big)$, $det\big(H_c(j\omega) + H_c^H(j\omega)\big) > 0$ for all $\omega$.

Consequently, transfer function matrix (S30) is WSPR (see the respective definition in **Useful mathematical concepts**). We therefore conclude that closed-loop system (S29) is asymptotically stable. To confirm this, we compute the eigenvalues of its dynamics matrix: $-1.5 \pm j0.87$ and $-0.5 \pm j0.87$ (they all have negative real parts).

Finally, note that in case we had R-Regulator with only one inhibitory reaction (either $Y_1 + Z_1 \xrightarrow{k_4} Z_1$ or $Y_2 + Z_2 \xrightarrow{k_3} Z_2$ ) we can immediately see from ODE model (S28) that one of the target species - $Y_2$ or $Y_1$ respectively - would go to infinity since the corresponding derivative would always be positive.

### References


[1] Domitilla Del Vecchio and Richard M Murray. *Biomolecular feedback systems*. Princeton University Press Princeton, NJ, 2015.

[2] Sigurd Skogestad and Ian Postlethwaite. *Multivariable feedback control: analysis and design*. Vol. 2. Citeseer, 2007.

[3] Bernard Brogliato et al. *Dissipative Systems Analysis and Control: Theory and Applications*. Springer, 2020.
